# TopicVI: A Knowledge-guided deep interpretable model for resolving context-specific gene programs

**DOI:** 10.64898/2026.04.08.717220

**Authors:** Guoxin Cai, Wenyi Zhao, Xunuo Zhu, Yitao Lin, Binbin Zhou, Ji Cao, Qiaojun He, Bo Yang, Xun Gu, Xushen Xiong, Zhan Zhou

## Abstract

Mechanistic insights from single-cell and spatial transcriptomics largely rely on cell clustering, differential expression analysis, and interpretation through prior biological knowledge. However, this approach is often limited by the reliance on curated biological priors that fail to capture context-specific gene programs, particularly in complex disease states. To address this gap, we introduce TopicVI, a deep interpretable model that integrates established biological knowledge with data-driven refinement to discover context-dependent gene programs in single-cell and spatial transcriptomic data. TopicVI jointly infers cell clusters and gene topics using optimal transport to flexibly align prior gene programs with observed data while permitting context-specific refinements. Comprehensive benchmarking demonstrates that TopicVI outperforms existing methods in biological conservation, batch correction, topic coherence, and rare cell identification. TopicVI effectively disentangles multiple sources of biological variation, such as separating anatomy-specific expression patterns from disease-associated signatures in spatial transcriptomics. Applying TopicVI to glioblastoma datasets, we identify gene topics related to cell cycle regulation and EGFR signaling that reveal convergent tumor states across distinct drug perturbations. By integrating prior knowledge with data-driven discovery, TopicVI enables identification of interpretable gene programs that illuminate biological processes and therapeutic mechanisms in complex transcriptomics data.

## Introduction

Single-cell RNA sequencing (scRNA-seq) has revolutionized our understanding of cellular heterogeneity and biological processes at unprecedented resolution ^1,2^. Spatial transcriptomics extends these capabilities but introduces unique analytical challenges. Spot-based technologies like 10x Visium capture transcriptomes from locations containing multiple cells (typically 5-20 per spot) ^3^, complicating both cell type deconvolution ^4,5^ and spatial domain identification ^6,7^. The standard analytical pipeline for both single-cell and spatial transcriptomics typically involves clustering followed by differential expression analysis to identify enriched biological functions and pathways ^8^. However, this conventional approach faces significant limitations when analyzing cells with subtle transcriptional differences or investigating perturbations that do not induce dramatic expression changes.

These challenges are particularly acute in disease contexts, where researchers seek to identify functionally altered cell subpopulations that may drive pathology or represent adaptive responses to disease conditions ^9,10^. In cancer research, characterizing immune cell expression profiles has become a critical focus. Many studies designate functionally altered cells as novel cell types ^11–13^, though this classification reflects functional rather than developmental divergence from traditional cell type definitions. These cells may retain core functional characteristics of their parent cell types while acquiring new functional properties or losing specific functions under pathological conditions. The scarcity of comprehensive datasets and corresponding annotations in such contexts poses additional challenges, particularly for supervised learning approaches ^14^.

To establish mappings between molecular phenotypes and disease states, researchers have developed various interpretable frameworks based on deep learning and machine learning models ^15^ (Supplementary Table 1). These approaches can be broadly categorized into two classes: *post-hoc* and *ante-hoc* explanations. Post-hoc explanations primarily assess the importance of individual features or genes by analyzing their contribution weights in model predictions. For instance, Shapley additive explanations (SHAP) ^16^ quantifies feature importance, where higher weights indicate greater influence on predictions. Ante-hoc explanations emphasize interpretable design within the model architecture itself. One mainstream ante-hoc approach constructs gene sets based on prior biological knowledge, incorporating gene programs annotated with established biological functions. For example, TOSICA ^17^ encodes gene expression into distinct biological pathways before processing through a transformer model, with downstream interpretation focusing on pathway-level activity. Another ante-hoc strategy builds inherently interpretable models where self-consistent interpretation emerges from the model architecture. Examples include non-negative matrix factorization (NMF), which decomposes expression matrices into feature modules and their sample loadings ^18–20^, and custom-designed interpretable components. Topic modeling, adapted from natural language processing, has been applied to RNA-seq data analysis ^21–23^. To model transcriptome heterogeneity, topic modeling treats genes as “words”, cells as “documents”, and biological functions as “topics”. Amortized latent Dirichlet allocation ^24^ integrates topic modeling within a variational autoencoder (VAE) framework to construct gene programs from cell embeddings. BALSAM ^21^ introduces spike-slab priors in the latent Dirichlet allocation (LDA) framework, assuming sparse gene weight distributions within programs, while Larch ^25^ further imposes hierarchical tree-like relationships between topics.

Despite these advances, existing solutions face a fundamental trade-off between biological interpretability and context-specific discovery. Prior-guided methods that rely strictly on established annotations ensure biological interpretability and leverage accumulated pathway knowledge, but they are constrained by the static nature of existing databases and cannot discover context-specific variations of known pathways. Conversely, purely *de novo* approaches enable identification of dataset-specific patterns but often produce gene programs that are difficult to interpret biologically, show limited consistency across studies. These limitations become particularly acute in complex analytical scenarios. In spatial transcriptomics, for instance, gene expression patterns simultaneously reflect tissue architecture, cellular composition within spots, and pathological states. Disentangling these intermingled signals—such as isolating disease-specific programs from anatomy-driven patterns—requires methods capable of targeted signal extraction while accounting for multiple biological factors. Current interpretable methods ^22,23^ focus primarily on gene program discovery but rarely enable context-specific modeling that controls for particular sources of variation. Similarly, in disease contexts and drug perturbations, distinguishing cell-state-specific programs from broader cell-type signatures requires flexible frameworks that can resolve the interpretability-discovery trade-off by treating prior knowledge as a guide rather than a constraint.

To address these challenges, we propose **TopicVI**, a deep topic model integrating VAE and NMF architectures. Motivated by the observation that cell clustering and identification of differentially expressed genes exhibit mutual influence, TopicVI jointly models cell clustering and gene topic discovery, yielding accurate results by leveraging interactions between cell clusters and gene topics. To distinguish cell populations with highly similar expression profiles, TopicVI incorporates prior gene programs (PGP) with known biological functions while permitting data-driven topic refinement. An optimal transport process aligns constructed gene topics with PGPs by transforming probabilities into distances between genes in PGPs and constructed topics. This approach ensures that gene topics remain consistent with both observed data and prior knowledge, enabling the identified topics to substantially overlap with prior information without being rigidly constrained. Additionally, grounding analysis in prior gene functions facilitates convenient functional and mechanistic interpretation. By integrating a novel optimal transport module for prior-guided topic refinement with a deep biclustering architecture that jointly models cell clusters and gene programs, TopicVI overcomes the historical limitation of decoupled cell and gene modeling. This innovation enables the model to disentangle subtle biological signals previously confounded—such as distinguishing functional states within transcriptionally similar cell populations or isolating disease-relevant programs from background variation. In the following results, TopicVI’s ability to identify context-specific gene programs—including drug-responsive signatures in glioblastoma and immune cell states in peripheral blood—demonstrates its potential for uncovering latent functional programs in complex tissue microenvironments where traditional clustering fails to resolve meaningful biological variation.

## Results

### Knowledge-Guided Topic Modeling for Interpreting Transcriptomic Variance

TopicVI is a deep topic modeling framework that jointly performs cell clustering and gene topic modeling by integrating prior biological knowledge with single-cell or spatial RNA sequencing data (Figure 1A). Unlike existing approaches that treat cell clustering and gene topic identification as independent processes, TopicVI exploits the mutual influence between these tasks to achieve more accurate cell representations and biologically interpretable gene topics. The model addresses key limitations of current methods by incorporating established biological knowledge while enabling data-driven refinement of gene programs.

**Figure 1.**
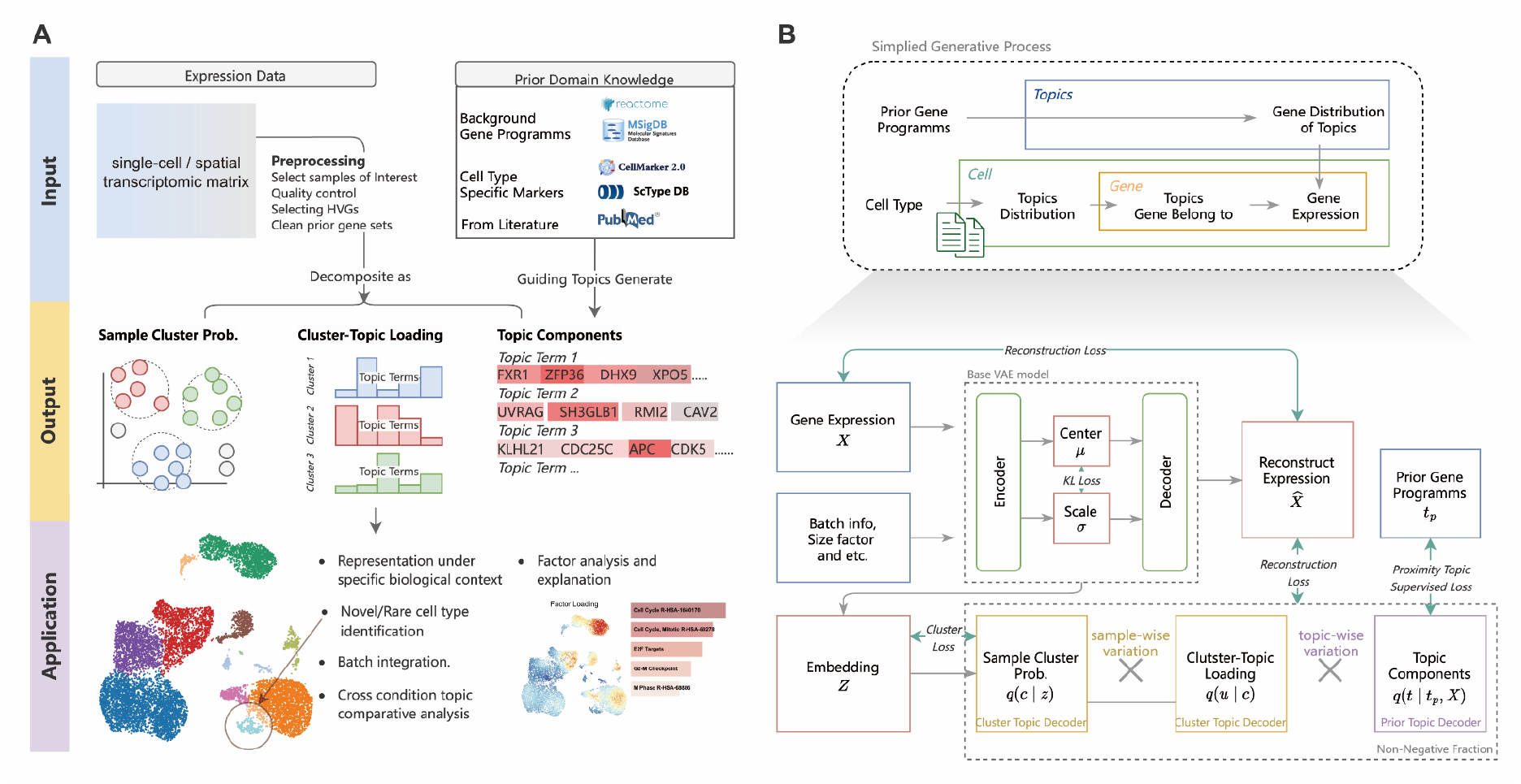
Topic modeling framework with variational inference incorporating prior biological knowledge. (A) Model architecture. TopicVI integrates gene expression data with prior biological knowledge to generate three output components: cell cluster assignment probabilities, cluster-topic loading matrix representing topic activity within each cluster, and topic-gene weight matrix defining gene weights within each topic. (B) Variational inference workflow. Upper panel: simplified generative model where prior gene programs inform topic-gene distributions and cell cluster assignments influence topic distributions. Lower panel: detailed TopicVI architecture combining VAE and NMF components. The VAE generates cell embeddings under clustering constraints, while prior gene programs guide topic formation via a proximity-based supervised loss. The original gene expression matrix is reconstructed through matrix multiplication of the three output components, constrained by biological priors. VAE, variational autoencoder; NMF, non-negative matrix factorization.

The TopicVI architecture comprises a cell representation module and a deep biclustering module that together generate three key output matrices: cell cluster assignment probabilities, cluster-topic loading matrix, and topic-gene weight matrix (Figure 1A). The cell representation module learns low-dimensional embeddings that capture cellular heterogeneity through a VAE architecture. The deep biclustering module operates on these embeddings to perform soft clustering via differentiable cluster centers positioned in the embedding space. A learnable parameter matrix maps clusters to topics, enabling each cluster to exhibit specific topic loading patterns. Integration of these three matrices reconstructs the expression data under biological consistency constraints.

To incorporate prior biological knowledge into topic construction and thereby ensuring both alignment with established biological pathways and discovery of novel functions, an optimal transport-based approach is applied (Figure 1B). Rather than rigidly constraining topics to match existing gene programs, the model performs semi-supervised topic learning by aligning constructed topics with prior gene programs through distance-based optimization (Methods). These distances derive from negative log-transformed probabilities of genes belonging to topics, enabling flexible alignment between data-driven discoveries and established biological knowledge. Consequently, this approach allows topics to maintain substantial overlap with prior information while accommodating novel functional relationships revealed by the data.

TopicVI outputs are interpreted through their relationships with prior biological knowledge and differential topic analysis. Cluster-specific topic enrichment is identified using statistical approaches analogous to differential gene expression analysis, typically employing t-tests to determine which topics are most active in each cluster. This dual validation approach—comparing topics to prior knowledge and identifying cluster-specific patterns—provides confidence in the biological relevance of discovered gene programs while revealing potential novel functional relationships that extend beyond existing annotations. The framework’s ability to simultaneously model cell clustering and gene topics while incorporating prior knowledge makes it particularly effective for complex single-cell datasets where traditional approaches may fail to capture the full spectrum of cellular heterogeneity and functional diversity.

### TopicVI Demonstrates Superior Performance Across HLCA Benchmarks

To comprehensively evaluate TopicVI’s performance, we conducted systematic benchmarking using the human lung cell atlas (HLCA) core data ^26^, which encompasses multiple levels of cell type annotation granularity. We constructed eight benchmark subsets via stratified sampling (see Supplementary Text) to simulate the complexity and heterogeneity characteristic of real-world single-cell RNA sequencing datasets. This benchmark comprehensively evaluated 13 methods across multiple dimensions, including quality of cell embeddings and gene topic components, with specific assessment of bio-conservation, batch correction, cluster accuracy, topic coherence, functional interpretability, and cell type differential expression (Figure 2A).

**Figure 2.**
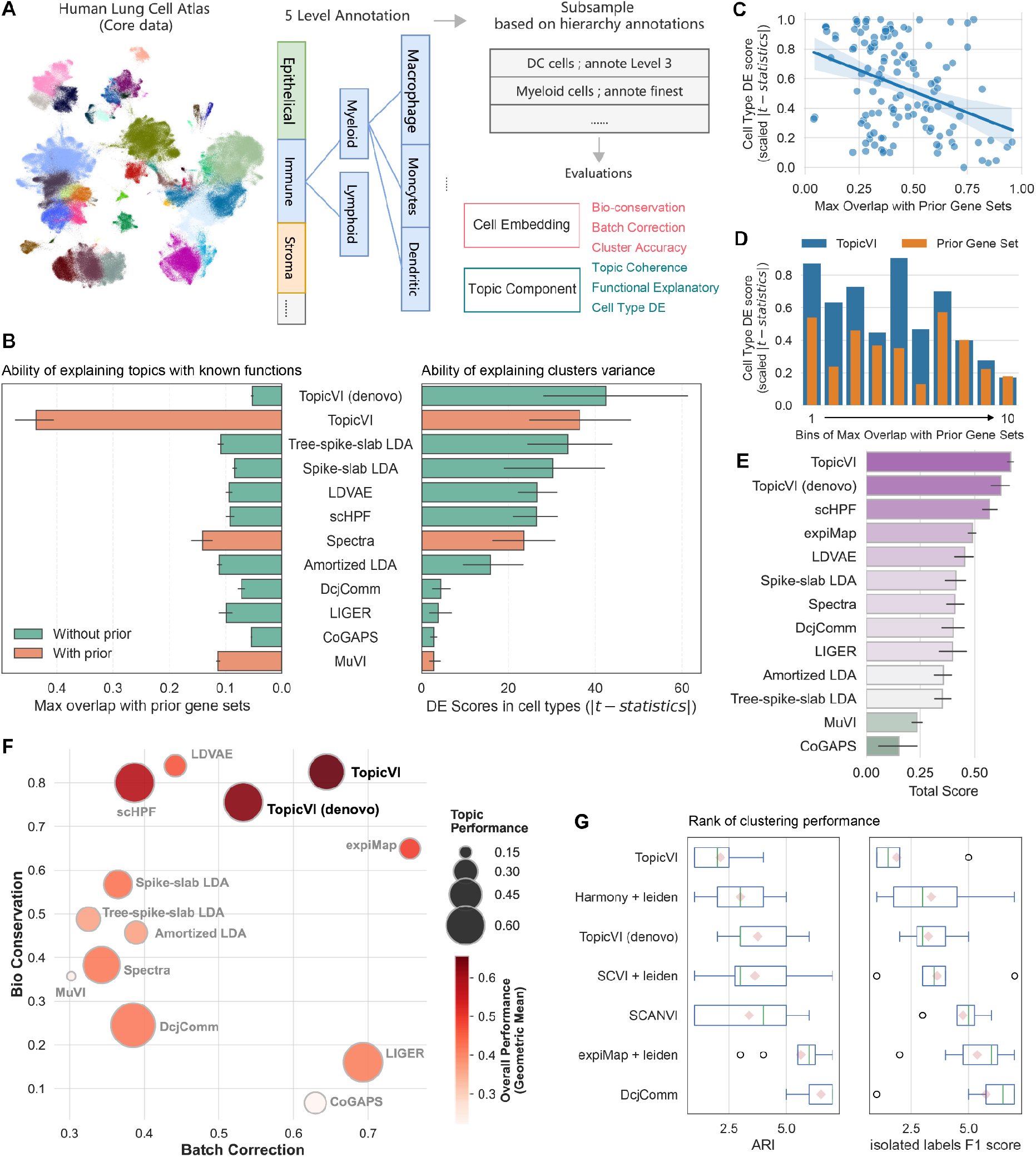
Comprehensive benchmarking of TopicVI using HLCA core dataset. (A) Evaluation framework. Eight datasets were subsampled at varying annotation granularities to enable systematic assessment of both cell embeddings and topic components. (B) Left panel: Gene program explanatory power measured by the Jaccard index, quantifying the maximum overlap between prior gene sets and model-derived gene programs. This evaluates how effectively each method incorporates prior biological knowledge. Right panel: Differential expression (DE) scores across cell types for each topic, defined as the maximum absolute t-statistic per topic across cell types, reflecting the ability to explain cluster-level variation. Bar colors indicate method categories based on prior knowledge usage. Error bars denote 95% confidence intervals. (C) Each point represents a topic, showing the correlation between its overlap with prior gene sets and its cell-type-specific DE score. (D) Topics from each dataset were grouped into ten bins based on their maximum overlap with prior gene sets. DE scores of topics and their corresponding prior gene sets were compared within each bin. (E) Total scores for 13 methods were computed by integrating three metrics: biological conservation, batch correction, and topic performance. (F) Integrated performance comparison across all evaluation metrics. Bubble positions represent performance in batch correction (x-axis) and biological conservation (y-axis), with bubble size reflecting topic performance scores. Bubble color represents the total score, which is the geometric mean of the three metrics. (G) Adjusted Rand Index (ARI) and isolated label F1 scores were used to evaluate clustering performance for both model-derived clusters and Leiden clustering applied to baseline cell embeddings. Box plots summarize performance ranks across the eight datasets (lower rank indicates better performance). Each box plot displays the interquartile range (IQR, box), the median (horizontal line), the mean (red diamond), and whiskers extending to 1.5×IQR. Outliers beyond this range are shown as individual points.

We first evaluated the interpretability of model-derived gene topics across two critical dimensions: biological function explanation and cell cluster variance explanation. To assess biological function explanation, we computed the maximum Jaccard index between model-inferred gene topics and predefined prior-knowledge gene programs. We found that models utilizing PGPs as input demonstrated overall greater consistency with prior-knowledge gene programs, while models without prior information showed relatively weak interpretability (Figure 2B). Notably, TopicVI with PGP input achieved the highest overlap, demonstrating superior ability to construct biologically meaningful gene programs by leveraging prior information. In contrast, other PGP-based models failed to effectively capture prior knowledge, exhibiting Jaccard indices comparable to *de novo* discovery methods. Cell cluster variance explained by gene topics was quantified using absolute t-statistics from differential expression analysis, analogous to standard differential expression analysis in single-cell transcriptomics (Methods). TopicVI demonstrated strong capability in constructing topics with high divergence across cell types, matching or exceeding all methods except TopicVI (*de novo*). This observation suggests a potential discrepancy between prior-knowledge functional gene topics and observed transcriptome data. To investigate whether this gap exists and how TopicVI addresses it, we examined the relationship between prior knowledge overlap and explanatory power. Overall, topics showing high overlap with prior knowledge exhibited lower cell type differential expression scores, suggesting a trade-off between these two axes (Figure 2C). We grouped all topics into ten bins based on their overlap with prior gene programs and compared their cell type differential expression scores with corresponding prior gene programs. Topics constructed by TopicVI showed largely stronger ability to explain cell type variance than their corresponding prior programs (Figure 2D). Collectively, while prior gene programs may inadequately explain data variance in specific contexts, our model successfully leverages prior knowledge to identify gene topics that better explain context-specific biological expression patterns.

To provide holistic performance evaluation, we further assessed three key metrics: biological signal preservation (bio-conservation), technical batch effect removal (batch correction), and coherence of generated gene programs (topic performance). Specifically, batch correction quantifies the extent to which batch effects are mitigated in cell embeddings; bio-conservation assesses alignment between cell embeddings and manual cell type annotations; and topic performance measures the divergence, similarity, and biological relevance of inferred gene topics (see Supplementary Text for details). We show that TopicVI achieves the best overall performance with an aggregate score of 0.67 (Figure 2E, Figure S1). In contrast, TopicVI without PGP supervision (TopicVI *de novo*) showed decreased performance in both batch correction and bio-conservation, although its topic performance score slightly improved (Figure S1, Table S3). This observation reveals that gene relationships predefined in PGPs display weaker correlations than *de novo* identified gene topics when analyzed in actual datasets; thus, integrating information derived from specific data with knowledge extracted from prior annotations enhances overall model performance beyond using predefined prior gene programs alone. Other topic models were biased towards either bio-conservation or batch correction while compromising the other. For example, scHPF ^27^ and LDVAE ^28^ emphasized biological factor representation such as cell type (achieving scores up to 0.8) but performed poorly in batch integration (scores at most 0.44), while LIGER conversely focused on batch correction. ExpiMap ^29^, which achieved similar overall performance to TopicVI, received a lower topic performance score (Figure 2F, aggregate score of 0.24), specifically showing poor gene program diversity and coherence. This may be attributed to its construction of gene programs without PGP guidance.

We next compared clustering performance between models that directly predict cell labels or clusters (TopicVI, scANVI, DcjComm) against those generating embeddings for downstream clustering (Harmony, ExpiMap) (Figure 2G, Table S4). We evaluated overall clustering accuracy using adjusted rand index (ARI) and the ability to identify rare cell populations using isolated label F1 score. Across all the datasets, TopicVI demonstrated the best performance compared to other commonly used methods. Notably, isolated label F1 score proved to be a more discriminative metric for evaluating clustering performance in datasets containing rare cell subtypes ^30^, as several methods performed well in ARI but poorly in identifying rare subtypes. Overall, extensive benchmarking on HLCA data demonstrates TopicVI’s capability in identifying meaningful gene topics, producing representative cell embeddings, and dissecting transcriptome heterogeneity across cell populations.

### TopicVI Resolves Immune Cell Subtypes in PBMC

Peripheral blood mononuclear cells (PBMCs) contain diverse immune cell populations whose functional states—such as activation, memory, and exhaustion—shift in response to stimuli. Expression differences between these states are often subtle, posing significant challenges for traditional methods to distinguish nuanced changes and identify fine-grained cell subpopulations. We applied TopicVI to dissect the complex cellular landscape of PBMCs, demonstrating how its interpretable gene programs can distinguish fine-grained cell subtypes and identify novel functional states. While higher clustering resolution may reveal finer substructures within cell embeddings—potentially corresponding to more granular cell subpopulations—an increased number of clusters does not necessarily imply improved biological interpretability but may instead represent forced sub-clustering lacking biological rationale. To account for the influence of clustering resolution on performance evaluation, we employed the Davies-Bouldin score to identify optimal resolution in subsequent analyses.

We applied TopicVI, scANVI, and scVI to the PBMC10k dataset (Figure 3A, Figure S1A). After performing Leiden clustering on cell embeddings, the Davies-Bouldin index was used to determine optimal clustering resolution. While TopicVI and scANVI achieved comparable normalized mutual information (NMI) scores (approximately 0.82), TopicVI achieved higher isolated label F1 scores than scANVI, and both significantly outperformed scVI. The results indicate that isolated label F1 score provides a more discriminating metric for evaluating performance on rare cell subtypes, especially in datasets with imbalanced cell populations. In detailed comparison, scANVI failed to resolve finer-grained cell subtypes, particularly within the myeloid lineage (gray circle in Figure 3A). In contrast, TopicVI effectively distinguished *FCGR3A*+ monocytes from *CD14*+ monocytes, dendritic cells (DCs), and megakaryocytes. These results together highlight the advantage of incorporating prior gene programs to resolve complex cellular hierarchies, particularly when rare or closely-related cell subtypes exist (Figure S2A).

**Figure 3.**
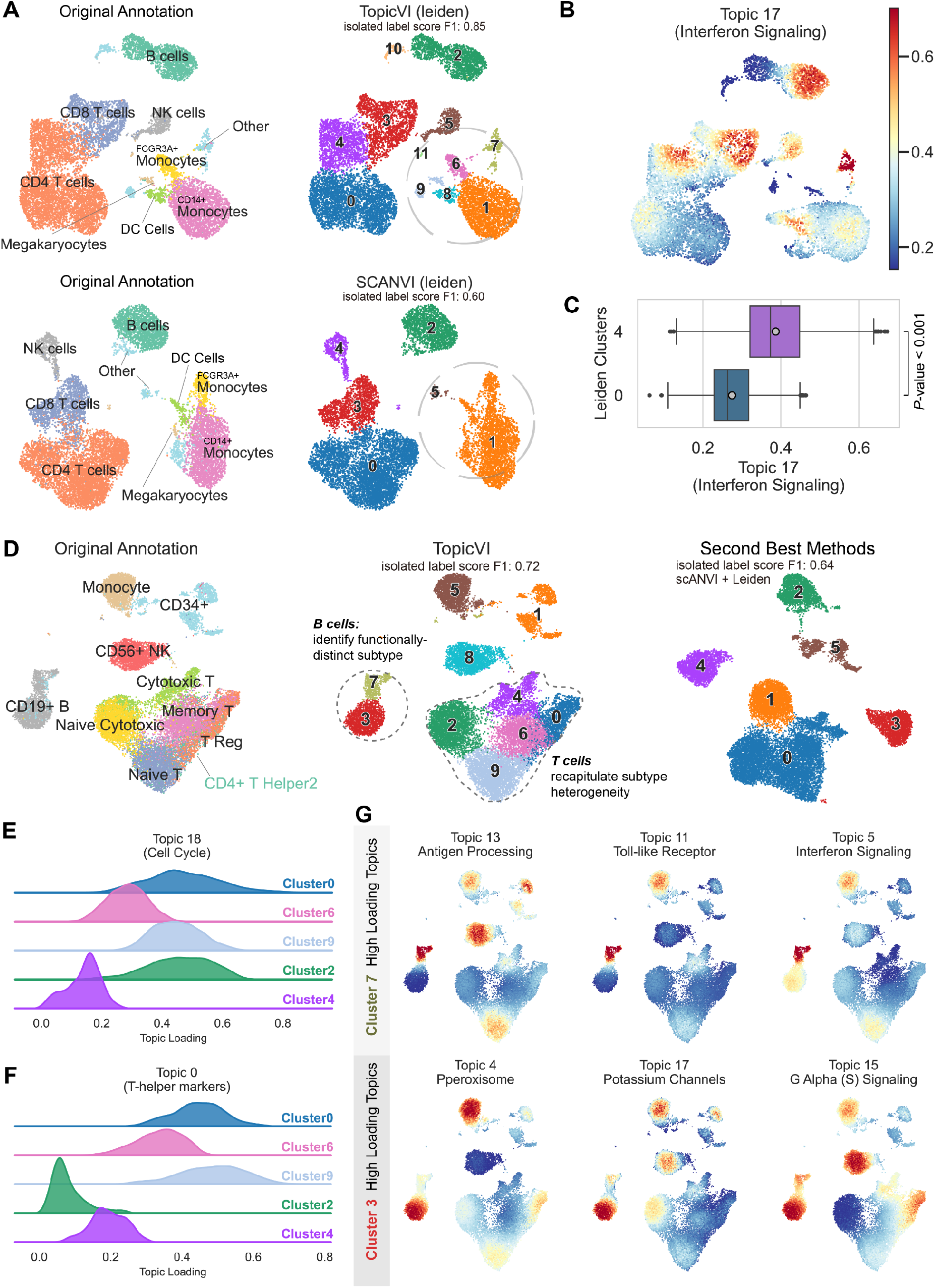
Cell type-specific topic analysis in peripheral blood mononuclear cell (PBMC) datasets. (A) UMAP visualizations of Leiden clustering applied to cell embeddings from TopicVI and scANVI on the PBMC10k dataset. TopicVI resolves myeloid cell subtypes with greater granularity than original annotations. (B) UMAP showing the loading distribution for Topic 17, annotated as the interferon signaling pathway. (C) Boxplot comparing Topic 17 loading between Cluster 4 and Cluster 0, showing significantly elevated values in Cluster 4 (Mann–Whitney U test, *P* < 0.001). (D) UMAPs illustrating TopicVI performance on the Zheng68k dataset, comparing TopicVI, the second-best method (Leiden clustering on scANVI embeddings), and manual annotations. (E-F) Violin plots showing topic loading distributions across T cell clusters for Topic 0 (T-helper cell markers, E) and Topic 18 (cell cycle-related genes, F). (G) Differential topics distinguishing B cell Cluster 7 from Cluster 3. Colors represent topic loading values, with red indicating higher and blue indicating lower values.

Furthermore, TopicVI subdivided *CD4*+ T cells into two distinct subpopulations (clusters 0 and 4). To assess whether our model captures biologically context-dependent transcriptome variance, we identified differentially activated gene topics between them (Figure S2B). Using a t-test with multiple testing correction on topic loadings, we found Topic 17 as the most significant distinguishing feature (Figure 3B-C). Based on input PGPs, we annotated this gene program as interferon signaling-related (Figure S2C). Therefore, based on Topic 17’s functional annotation, we classified cluster 4 as activated *CD4*+ T cells differentiating into T-helper cells, while cluster 0 corresponds to naive *CD4*+ T cells. Notably, existing research highlights the critical role of interferon in *CD4*+ T cell differentiation and maturation ^31,32^, where it promotes their activation and differentiation into T-helper cells. We also investigated an undefined population (labeled “Others” in the original study), which TopicVI confidently separated into clusters 7 and 9. Functionally, cluster 7 exhibited high loadings for Topic 11 (apoptosis-related), Topic 0 (myeloid cell markers), and Topic 4 (interleukin signaling) (Figure S2D). These functional signatures indicated that cluster 7 represents macrophage or neutrophil populations, which is a previously undefined cell cluster ^13,33^.

To evaluate TopicVI’s ability in resolving higher-resolution immune profiles, we further applied TopicVI to the Zheng68k PBMC dataset ^34,35^. This dataset was pretreated with fluorescence-activated cell sorting (FACS), which provided more granular partitioning based on cell surface proteins than the manually annotated PBMC10k dataset. TopicVI demonstrated superior clustering performance in dissecting complex cell heterogeneity, as shown by the higher isolated label F1 score (Figure 3D, Figure S3A). In contrast, scANVI, which uses a seed-labeling strategy with marker genes from other datasets, produced output that differed significantly from reference annotations (Figure S3C). This discrepancy likely arises because cell type marker genes are not always robust across datasets, especially when influenced by batch effects or different biological contexts. TopicVI, however, adjusts its gene programs based on input data, thereby achieving more contextually robust results. Comprehensive comparative analysis further confirmed TopicVI’s effectiveness in distinguishing various T cell subtypes, which other methods struggled to achieve (Figure S3B-F). To explore T cell subtypes, we examined two characteristic topics: Topic 0 (helper T cell markers) and Topic 18 (cell cycle regulation) (Figure 3E-F). Among T cell subpopulations, clusters 2 and 4 exhibited lower Topic 0 loadings, consistent with their annotation as cytotoxic T cells. Topic 0 loading patterns in remaining subpopulations (memory T cells, regulatory T cells, naive T cells, and Th2 cells) were consistent with manual annotations. Notably, Topic 18 showed low loadings exclusively in cytotoxic T cells, aligning with their terminally differentiated state. This concordance between topic loading patterns and manual annotations confirms the reliability of our analytical approach, demonstrating that TopicVI resolves cell subtypes with higher fidelity, especially for diverse functional states of T cells (clusters 0, 2, 4, 6, 9).

Topic modeling by TopicVI also enables functional characterization of transcriptionally similar cell subtypes. TopicVI clustering identified two B cell clusters exhibiting different biological functions represented by distinct topic signatures (Figure 3G). Cluster 7 showed significantly elevated loadings on topics related to antigen presentation, Toll-like receptor pathways, and interferon signaling. In contrast, high-loading topics for cluster 3 were associated with fundamental biological processes, such as sodium ion channel regulation and G-protein coupled signal transduction. This differential pattern suggests that cluster 7 represents immunologically active B cells while cluster 3 represents resting B cells.

### Supervised Topic Modeling Disentangles Confounding Biological Signals in Human Brain Spatial Transcriptomics

A fundamental challenge in transcriptomics is that gene expression is often influenced by multiple biological factors simultaneously. For instance, both anatomical location (e.g., cortical layer) and pathological condition can drive expression changes, making it difficult to isolate the signal of interest from confounding variables. While spatial transcriptomics incorporates spatial information to improve domain detection ^3,6^, disentangling overlapping biological signals remains challenging. Notably, TopicVI can be used to address this limitation by adaptation to a supervised learning mode. This approach uses known annotations to guide the model, explicitly constructing gene programs associated with the target variable of interest. TopicVI incorporates a focal loss term into its objective function ^36^ (Supplementary Text), which proves particularly effective for handling class imbalance and noisy biological labels.

We applied this strategy to a spatial transcriptomics dataset of human cerebral cortex, which included annotations for both cortical layers and disease diagnosis. We first assessed whether supervised learning produces cell embeddings that faithfully represent training labels. The TopicVI model trained on anatomy layer labels was compared to unsupervised TopicVI, scVI, and Spectra with cell type awareness. Using the unsupervised mode of TopicVI and other methods, the resulting cell embeddings were influenced by both layer identity and pathological state, producing a convoluted representation (Figure S4A). In contrast, supervised mode produced a UMAP visualization with remarkably clear, layer-ordered structure reflecting known cortical anatomy, effectively separating layer-specific signals from confounding disease signals (Figure 4A).

**Figure 4.**
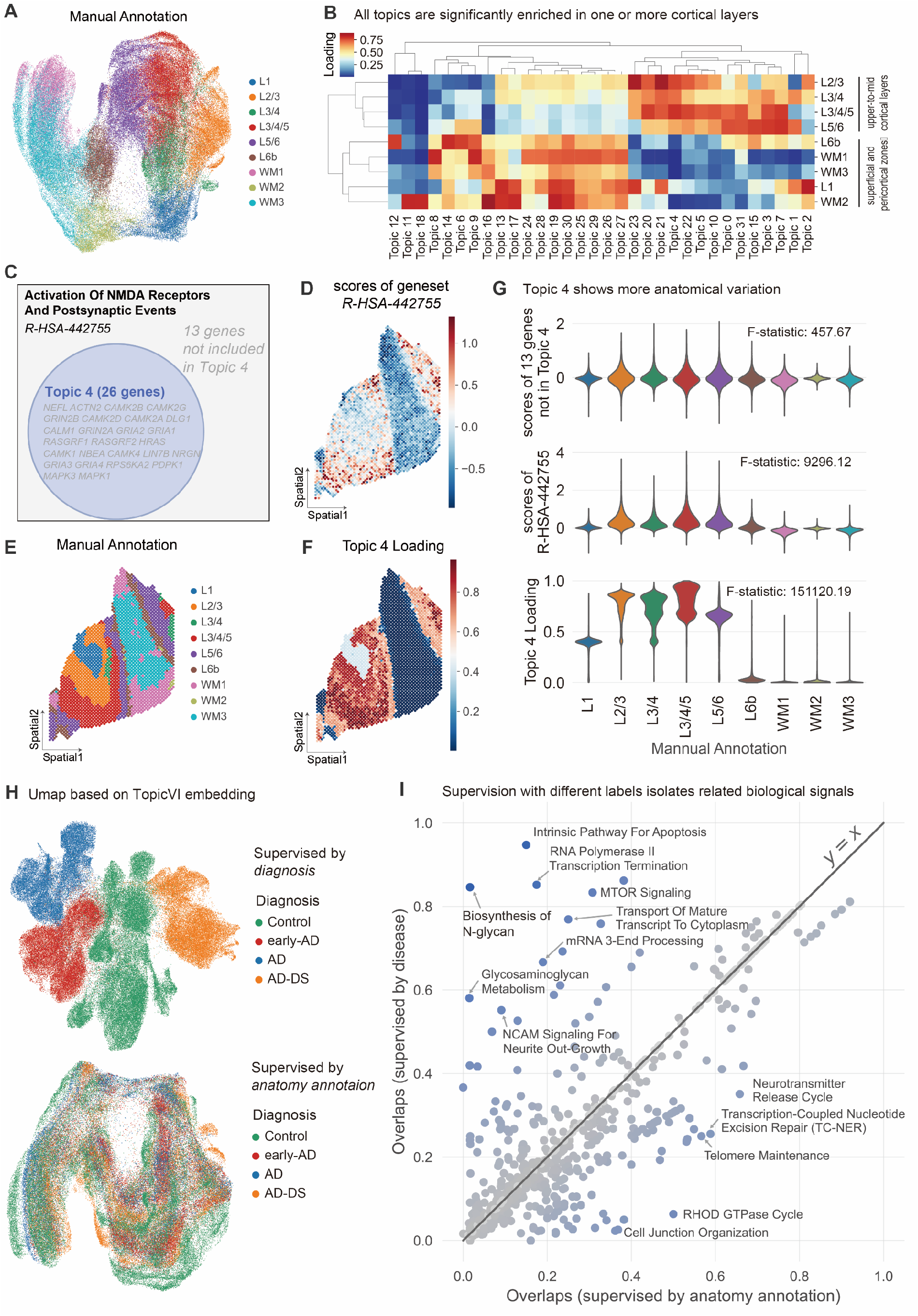
Supervised topic modeling isolates biological signals in human brain cortex. (A) UMAP visualization of cell embeddings colored by manual cell type annotations. (B) Layer-specific topic activity across cortical layers. Each topic shows significant differential loading across layers (Student’s *t*-test). The heatmap displays average topic loading per layer, with rows and columns clustered by similarity. (C) Topic 4 shows substantial overlap with Reactome pathway R-HSA-442755, associated with NMDA receptor activation and post-synaptic signaling. (D-F) Spatial distribution patterns of R-HSA-442755 pathway scores (D), manual annotations (E), and Topic 4 loading (F). (G) Violin plots comparing layer specificity of Topic 4, the corresponding Reactome pathway, and 13 excluded genes. Topic 4 exhibits markedly higher cluster-specific resolution (F-statistic: 151,120.19 vs. 9,296.12). (H) Supervised learning enables targeted modeling of specific biological variables while controlling for confounding factors. UMAPs show TopicVI results when supervised by either anatomical annotations or disease diagnosis. (I) Supervision with distinct labels isolates different biological signals. The scatter plot compares Jaccard index overlaps with prior gene sets under two training regimes: supervision by diagnosis (y-axis) and by anatomy (x-axis). Each point represents a prior gene set used in TopicVI; point color reflects the magnitude of difference between the two training conditions. Selected gene sets with large differences are highlighted with labels and arrows.

Crucially, anatomically-supervised topics were highly specific to the anatomical labels used for guidance. We found that each topic was significantly differentially active in at least one specific cortical layer (Figure 4B), confirming that the model successfully learned distinct layer-specific gene programs. To examine the biological relevance of these programs, we focused on Topic 4, which showed high activity in upper-to-mid cortical layers (L2/3 to L5/6) (Figure S4B). Functional analysis revealed that Topic 4 was highly associated with NMDA receptor activation and postsynaptic signaling, derived from Reactome pathway “R-HSA-442755” (Figure 4C). Notably, TopicVI did not simply replicate prior knowledge. Instead, it performed data-driven pathway refinement: Topic 4 contained only 26 of the 39 (67%) genes from the original Reactome pathway, selectively excluding 13 genes from the canonical pathway. This data-driven refinement was functionally relevant. When we compared the layer-discriminating ability of Topic 4 against the original pathway, Topic 4 loading scores provided much clearer distinction between cortical layers (Figure S4C-D). This was further validated spatially, where Topic 4 activity maps mirrored manual layer annotations with high fidelity, appearing sharper and less noisy than activity maps of the full Reactome gene set (Figure 4D-F, Figure S5). Using F-statistic as a metric, we demonstrate that exclusion of 13 genes substantially improved layer discrimination (F-stat = 457.67 for excluded genes, one-way ANOVA test) compared to the complete gene program (F-stat = 9,296.12), while Topic 4 loading (F-stat = 151,120.19, *P* < 0.0001) significantly distinguished different anatomical layers (Figure 4G). This demonstrates TopicVI’s ability to refine prior knowledge to better fit observed data, improving biological specificity and revealing potential gene function changes under specific biological contexts.

We then compared model training with two different label types: anatomical layers and disease diagnosis. Comparing UMAP visualizations, disease diagnosis-supervised training separated cells by disease state while treating anatomical layers as confounding factors (Figure 4H). We hypothesized that topics derived from the two models would reflect biological processes related to their respective training labels. By examining maximum overlap with prior gene programs in both training strategies, we compared the alignment of each prior between topics. While the priors showed globally similar overlap between the two labels, several topics exhibited pronounced discrepancies (Figure 4I). Topics with larger overlap when supervised by anatomical layers included fundamental biological functions such as neurotransmitter release cycles. In contrast, gene programs including apoptosis ^37^, mTOR pathway ^38^, and biosynthesis of N-glycans ^39,40^ are established as Alzheimer’s disease-related, showing greater enrichment in disease-supervised topics (Table S5).

In conclusion, supervised TopicVI provides a powerful method for disentangling complex biological signals and discovering highly relevant, data-driven gene programs. This capability enables more accurate and nuanced interpretation of underlying biological processes and is broadly applicable to other spatial and single-cell datasets where confounding factors or prior annotations exist.

### TopicVI-Derived Topics Reveal Survival Associations and Drug Sensitivity Patterns in Glioblastoma

Having extensively demonstrated TopicVI’s ability to resolve distinct subtypes in a functionally interpretable manner, we next applied it to investigate drug perturbation and treatment responses using a glioblastoma (GBM) dataset comprising tissue slices exposed to drugs ^41^. We first performed cell type annotation for TopicVI clustering results (Figure 5A-B, Figure S6A), using published cell markers and malignancy scores derived from the expression ratio of chromosomes 7 and 10, which reflect copy number alterations characteristic of GBM (chr 7 amplification and chr 10 deletion).

**Figure 5.**
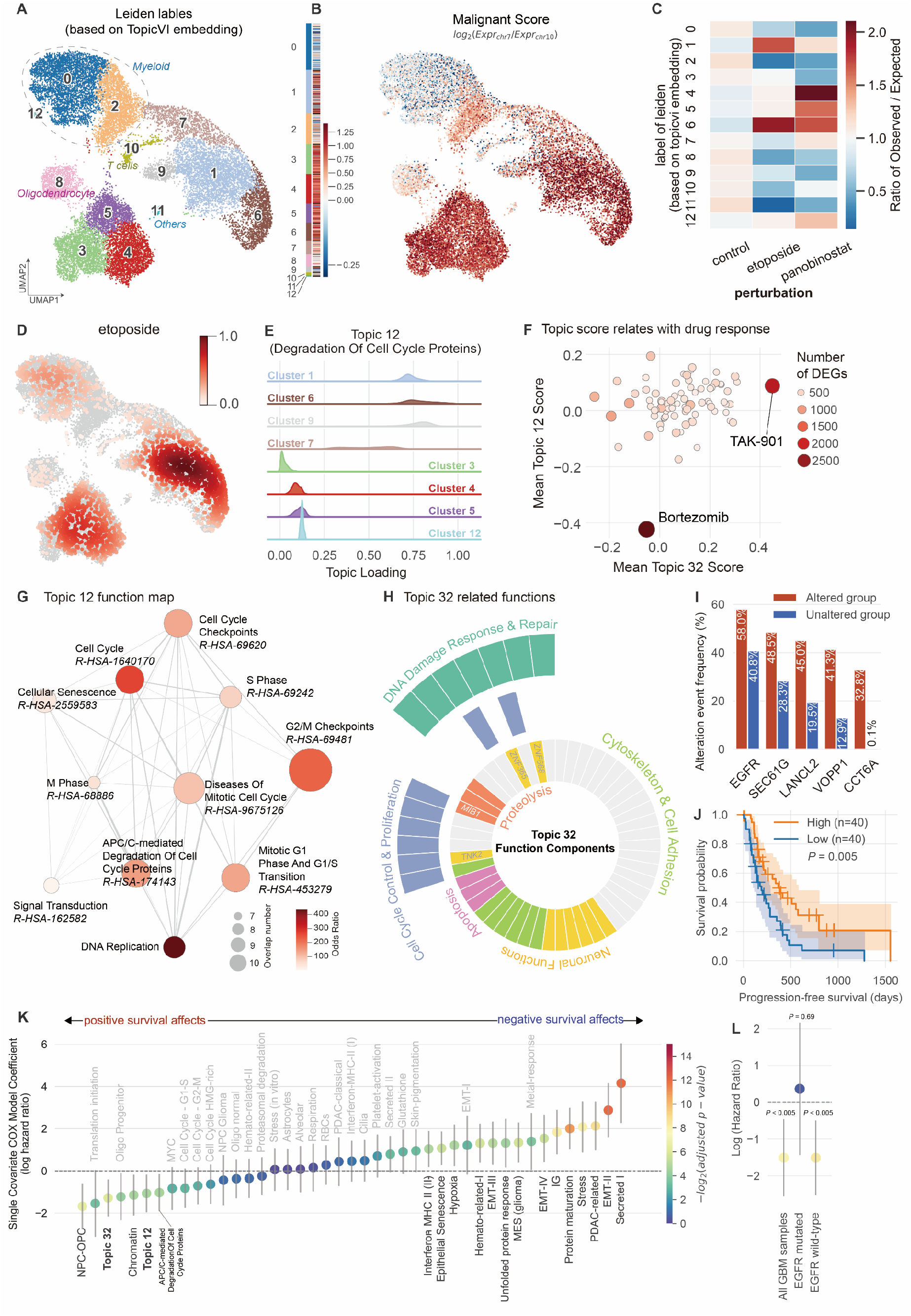
TopicVI reveals drug perturbation effects in glioblastoma tumor cells. (A) UMAP visualization of cell clusters based on TopicVI embeddings, annotated with Leiden clustering labels. (B) Malignant score quantified by log-transformed ratio of average expression between chromosome 7 and chromosome 10. (C) Enrichment analysis showing the ratio of observed to expected perturbation cells across clusters. (D) Relative cell density under etoposide treatment conditions. (E) Topic 12 loading distribution (associated with degradation of cell cycle proteins) across clusters. Clusters 1, 6, and 9 exhibit significantly higher loading compared to other groups. (F) Topic scores in relation to drug response in the A-172 cell line. Each point represents a drug, with point size and color indicating the number of DE genes compared to the DMSO control. TAK-901 and bortezomib show over 2,000 DE genes and exhibit the highest Topic 32 scores and lowest Topic 12 scores. (G) Functional enrichment network for Topic 12. Node size represents gene overlap count, node color indicates odds ratio, and edges denote functional similarity (Jaccard index). (H) Gene functional analysis for Topic 32. Gene descriptions were manually categorized; TNK2, MIB1, ZNF365, and ZNF688 are highlighted due to known associations with glioblastoma (GBM). (I) Frequency of EGFR alteration events and co-amplified genes, grouped by whether genes in Topic 32 are altered or not in 8 independent studies. (J) TCGA survival analysis stratified by Topic 32 scores. Patients in the top quartile (highest Topic 32 scores) show significantly longer progression-free survival than those in the bottom quartile (log-rank test, *P* = 0.005). (K) Single-covariate Cox model showing hazard ratios (HR) for Topic 32, Topic 12, and meta cancer programs from Gavish et al. Topics 32 and 12 demonstrate significantly stronger positive survival associations. The grey line represents the 95% confidence interval of HR. (L) Stratified Cox proportional hazards regression showing the association between Topic 32 score and progression-free survival in all GBM patients, EGFR-mutant patients and EGFR wild-type patients.

TopicVI separated malignant cells into distinct subpopulations, highlighting substantial functional heterogeneity within the tumor. We subsequently applied topic modeling to uncover the molecular basis underlying this heterogeneity. By calculating the ratio of observed to expected cell frequencies per cluster, we identified significant enrichment of etoposide-treated cells in clusters 1 and 6 (Figure 5C-D), which was not reported in the original research. To understand the basis of this enrichment, we examined cluster-specific topics and found that Topics 12, 21, and 23 show strong activity in these etoposide-responsive clusters (Figure S6B). These topics correspond to key pathways including APC/C-mediated degradation of cell cycle proteins, regulation of *MECP2* expression, and the RND GTPase cycle (Figure 5E, Figure S6C). which are highly relevant to GBM biology and etoposide mechanism of action. For instance, MECP2 regulates epithelial-to-mesenchymal transition in GBM ^42^, RND GTPase controls p53 function ^43^, and the anaphase-promoting complex/cyclosome (APC/C)—a large E3 ubiquitin ligase crucial for cell cycle regulation—exhibits documented synthetic lethal interactions with etoposide and the APC/C-CDH1 complex ^44^. Based on these functional annotations and enrichment patterns, we defined clusters 1, 6, 7, and 9 as drug-responsive tumor cell clusters.

To further confirm that these topics are related to drug response, we utilized the A-172 cell line as an external validation, which exhibited the second highest Topic 32 score among 33 GBM cell lines in the CCLE database (Figure S7A) and possessed systematic drug perturbation single-cell data in the Tahoe-100M database. By comparing number of differentially expressed genes between drug-perturbed cells and control groups, we found that two drugs with the largest differential expression profiles exhibited the highest and lowest topic scores for Topic 32 and Topic 12, respectively (Figure 5F).

We observed that treatments with bortezomib and etoposide—despite initiating cellular stress through distinct primary targets—resulted in a convergent phenotypic state characterized by Topic 12 activity. The down-regulation of Topic 12 (cell cycle progression) under bortezomib treatment aligns with its mechanism of inducing G2/M arrest (Figure 5G, Figure S7B). This convergence suggests a “dual-hit” vulnerability: the shared transcriptional bottleneck implies that combining these agents could prevent GBM cells from bypassing cell cycle checkpoints, a synergy previously observed in myeloma ^45,46^, leukemia ^45^, and prostate cancer ^47^ cell lines.

Notably, Topic 32, a gene program constructed *de novo* by TopicVI, was also strongly associated with etoposide response. As this topic exhibited minimal overlap with our prior knowledge annotations, we performed manual functional annotation by collecting gene summaries from multiple databases (Figure 5H, Table S6). Our analysis revealed that Topic 32 comprises genes involved in multiple cancer-related processes, including cell cycle control, DNA damage response, and apoptosis. It also contains several established drug targets or known biomarkers, such as *TNK2*, a target of lorlatinib for non-small cell lung cancer ^48^, dasatinib for leukemia ^49^, and *MIB1*, a known prognostic marker for GBM ^50^. We have also found that genes in the Topic 32 show high co-mutation with genes in the *EGFR* pathway (Figure 5I), implicating that this topic may be relevant to a key malignant biological process as the *EGFR* is the commonly acknowledged cancer driver. This analysis validates the biological relevance of Topic 32 in characterizing drug-perturbed GBM cells and showcases TopicVI’s ability to discover novel, treatment-relevant gene programs directly from data.

To confirm the clinical relevance of the identified topics, we performed survival analysis using bulk RNA-seq data from the cancer genome atlas (TCGA). We calculated a Topic 32 score for each patient in the TCGA-GBM cohort and stratified them into high (top 25%) and low (bottom 25%) expression groups. Patients with high Topic 32 scores exhibited significantly longer progression-free survival compared to those with low scores (log-rank test, *P* = 0.005, Figure 5J), similar to patterns observed for known cancer meta-programs (Figure S7C). We also conducted single-covariate Cox regression comparing these two topics with cancer meta-programs from Gavish et al. ^51^. Both Topic 32 and Topic 12 demonstrated significant positive survival effects (Figure 5K). Notably, Topic 32 showed a strong positive effect across all cancer meta-programs except neural progenitor cells with oligodendrocyte progenitors (NPC-OPC).

As the genes in Topic 32 showed genomic co-occurrence with the *EGFR* pathway, we next asked whether the prognostic value of this topic is dependent on *EGFR* mutation status. Intriguingly, stratified Cox models revealed that the protective effect was confined to *EGFR*-wild-type but not *EGFR*-mutated tumors (Figure 5L, Table S7). The induction of the program by both genotoxic and mitotic agents suggests that it represents a generalizable stress-response module. Tumors capable of mounting this response may show higher tolerance on therapy-induced damage, consistent with the improved survival observed in *EGFR*-wild-type patients. Although direct evidence that *EGFR*-mutant GBM is specifically resistant to etoposide is limited, EGFR alterations are broadly associated with treatment resistance, transcriptional plasticity, and therapy-tolerant states ^52–54^. In this context, our findings support a model in which EGFR-driven oncogenic signaling overrides or saturates the beneficial effects of drug-induced stress-response pathways, explaining why the program is prognostic only in EGFR-wild-type tumors.

These results demonstrate that TopicVI can decompose complex drug responses into distinct, interpretable gene programs, facilitating identification of mechanistic similarities between drugs and supporting applications in drug repurposing and combination therapy design.

## Discussion

In this study, we developed TopicVI, a data- and knowledge-driven model that identifies interpretable gene programs to elucidate cellular heterogeneity. Our approach leverages prior biological knowledge while using data-driven discovery to identify context-specific modifications to established knowledge. By building on a VAE framework and incorporating concepts from LDA and NMF, TopicVI synergistically optimizes three interdependent tasks: deep cellular embedding, cell clustering, and prior-guided topic modeling. To guide gene topic construction from prior knowledge, we employed an optimal transport formulation solved efficiently using Sinkhorn entropy regularization. The inherent flexibility in solving optimal transport problems serves as beneficial regularization, encouraging resulting topics to better conform to the observed data distribution.

Unlike traditional topic models such as ProdLDA ^55^, which assume Dirichlet priors for topics and multinomial distributions for words, TopicVI is specifically designed for the unique characteristics of single-cell transcriptomics data. While some methods attempt to adapt multinomial distributions for scRNA-seq data ^21^, TopicVI directly addresses key technical challenges—such as dropout events and gene expression inflation—by employing a zero-inflated negative binomial (ZINB) distribution, a standard modeling choice in the field. Although conceptually built upon the LDA framework, TopicVI’s flexible architecture supports alternative data distributions, such as normal distributions, making it suitable for modeling normalized expression in pseudo-bulk data and broadly applicable to other omics modalities.

A fundamental challenge in scRNA-seq analysis is reconciling generalizable biological knowledge with dataset-specific features. Prior gene programs establish robust, generalizable cell markers across studies, yet cell-state-defining genes are often dataset-specific. Bridging this gap requires models that unify prior knowledge with data-driven signals—a complementarity realized through TopicVI’s architecture. Our benchmarking results reflect this balance: the full TopicVI model outperformed the *de novo* version (without priors) on bio-conservation and batch correction, demonstrating the generalizability conferred by incorporating priors. Conversely, the *de novo* method showed higher topic diversity, as it was not constrained by inherent overlaps present in curated prior gene programs. However, the full TopicVI model excelled in topic coherence, as connections within prior gene programs are typically stronger and more biologically established.

TopicVI was also applied to drug perturbation datasets to identify cellular responses to treatment. Virtual cell modeling has emerged as a central challenge ^56^ for predicting potential cellular responses to drugs, gene knockouts, and environmental perturbations, which may facilitate drug repurposing and drug target identification. However, the primary challenge lies in out-of-distribution generation and few-shot prediction capabilities. Topic modeling offers potential solutions to these limitations and provides a foundation for virtual cell modeling through two key features: integration of prior biological knowledge enhances interpretability and mechanistic understanding, while the VAE-based framework enables robust data generation. By mapping perturbations to distinct topics, topic modeling may leverage the inherent robustness of biological mechanisms to potentially overcome out-of-distribution generation challenges. This topic-based representation of cellular responses provides a biologically grounded approach to understanding how perturbations affect cellular states, offering a pathway toward more reliable virtual cell models that can generalize beyond training data distributions.

In summary, TopicVI provides a powerful framework for constructing and discovering gene programs by integrating prior knowledge with data from specific biological contexts. Through comprehensive benchmarking and application to diverse datasets, TopicVI has demonstrated superior performance in explaining cell subtype differences, identifying functional state changes, and revealing cellular responses to perturbations. This work offers a new and effective strategy for enhancing biological interpretation of complex transcriptomic data.

## Methods

### Preprocessing for RNA-seq Data

We followed the best practices for scRNA-seq data analysis as described by Lukas et al. ^8^. First, uniqueness of gene and cell identifiers was verified, and key biomarkers including mitochondrial genes, ribosomal genes, and hemoglobin genes were labeled for subsequent quality control metrics. Multiple quality control indicators were calculated at the cellular level, including the total UMI count, the number of detected genes, the percentage of highly expressed genes, and the mitochondrial gene expression ratio.

Quality control employed the median absolute deviation (MAD) as the core assessment metric, calculated as follows:

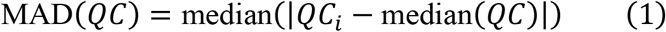

where *QC* represents the quality control metric being assessed, and median denotes the median of values. A dual filtering mechanism was established based on this metric: a 5-fold MAD threshold was applied to total log UMI, log genes detected, and the percentage of highly expressed genes to filter out low-quality cells, while a 3-fold MAD threshold or a composite criterion of an absolute threshold exceeding 8% was imposed on mitochondrial gene expression to effectively exclude cells with aberrant mitochondrial activity.

Cells passing quality control screening proceeded to the normalization process, where raw counts were normalized to a baseline level of 10^4^ total expression, followed by logarithmic transformation to mitigate the influence of highly expressed genes. For scRNA-seq data, we used `scanpy.pp.highly_variable_genes` (Seurat_v3 algorithm) ^57,58^ to identify 2,000 highly variable genes (HVGs). For spatial transcriptome data, we employed the “Spanve” method ^59^ to identify spatially variable genes (SVGs). For multiple slice analysis, we selected the union of SVGs from each slice.

### Prior Knowledge Collection and Processing

TopicVI requires prior knowledge in the format of gene sets collected from databases proven to be biologically meaningful. Prior knowledge includes two types: background gene knowledge and cell type markers. Cell marker genes characterize hallmark expression patterns of specific cell types or their functional states, while cell function-related genes serve as background gene programs incorporated into the model to maintain characterization of underlying biological functions.

Background gene sets involved in this research contain gene sets from Reactome DB ^60^ and MSigDB Hallmark ^61^. For cell type markers, we selected different gene annotation sources based on biological and pathological context. For scRNA-seq analysis of PBMC data, CellMarker DB V2.0 ^62^ and ScType annotation ^63^ were used based on original dataset annotations. For GBM tissue analysis, we incorporated meta-programs of malignant cells constructed by Gavish et al. ^51^. For human cortex Visium data analysis, we filtered cell markers specific to brain tissue from ScType annotation.

Before input into the model, collected prior gene programs (PGPs) were rigorously screened: first, each PGP was filtered to retain only the intersection with data variables; subsequently, entries with fewer than five genes in background gene programs and fewer than three genes in cell type marker programs were removed.

### TopicVI Model Architecture and Training Procedure

TopicVI is implemented using the M1+M2 framework ^64^ to achieve variational inference for gene topic identification and sample clustering. The framework comprises two main modules: a cell embedding module and a deep biclustering module.

The cell embedding module is based on the scVI model ^24^ and is primarily responsible for batch effect correction and expression modeling of single-cell data to construct cell embedding representations. The model adopts a standard encoder-decoder architecture. The encoder contains two layers of 128-dimensional fully connected networks with batch normalization and layer normalization, ultimately outputting 32-dimensional cell embedding representations in latent space. The decoder employs linear layers with softmax activation function, achieving expression reconstruction through construction of four parameters of the ZINB distribution using reparameterization tricks:

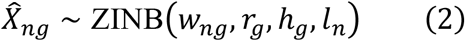

where parameter meanings are as follows: *w*_*ng*_: distribution mean; *r*_*g*_: distribution dispersion; *h*_*g*_: zero-inflation probability; *l*_*n*_: library scale factor. These parameters are regressed through neural network decoders. To guide training of the cell embedding module, an evidence lower bound (ELBO) loss constrains model parameters (see Supplementary Text for details). In training the TopicVI model, we first pretrain a scVI model, then transfer parameters to the cell embedding module to reduce running time and computational cost.

To achieve integration of gene topic modeling and cell clustering, the deep biclustering module describes the gene expression mean *w*_*ng*_ using three components:

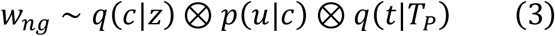

where: *q*(*c*|*z*) represents cell cluster assignment probability after deep clustering; *p*(*u*|*c*) represents gene-topic probability distribution corresponding to cell clusters; *q*(*t*|*T*_*P*_) represents gene weights within gene topics under guidance of prior topic *T*_*P*_.

To ensure robustness across high and low expression ranges, we do not directly reconstruct expression through multiplication but instead model gene expression proportions and construct loss functions using maximum likelihood principles:

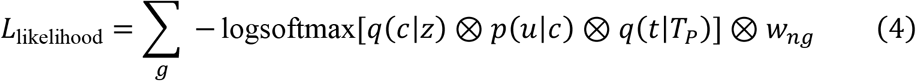

where ⊗ represents tensor product operation. Among the above parameters, *q*(*c*|*z*) is provided by the deep clustering process, *q*(*t*|*T*_*P*_) is provided by the topic optimization transfer process, while *p*(*u*|*c*) are differentiable model parameters.

### Semi-Supervised Gene Topic Modeling

The gene-topic assignment probabilities 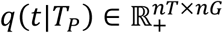 are generated through a semi-supervised learning process, where *nT* represents the number of topics and *nG* is the number of genes. To enforce sparsity in *q*(*t*|*T*_*P*_) (as topics are expected to contain only a small subset of all genes), we have *q*(*t*|*T*_*P*_) = softmax(*qT*), where *qT* is a model parameter.

When training the TopicVI model, an optimal transport algorithm aligns topics *T* = {*t*_1_, *t*_2_, *t*_3_, …} with priors *T*_*P*_. In the optimal transport process, the distance between topic *T* and prior *T*_*P*_ is defined as the average distance of genes in *T*_*P*_, and the distance between genes and topic *T* is defined as the negative log-probability of genes belonging to the topic:

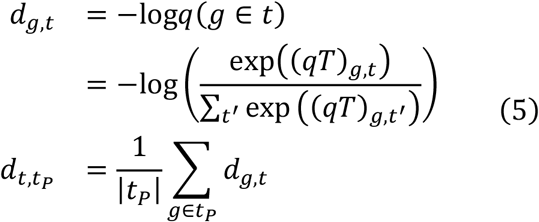

The optimal transport loss is then formulated as:

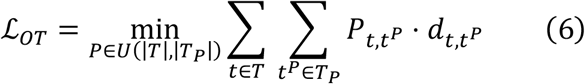

where *P* ∈ *U* represents the transport plan. Existing research ^65^ has demonstrated that optimal transport problems based on log-probability transformations are equivalent to solving semi-supervised problems.

To improve computational efficiency, we employ Jean Feydy’s annealing algorithm ^66^ to calculate symmetric Sinkhorn divergence for solving optimal transport problems. Sinkhorn divergence ^67^ regularizes transport plans by adding entropy penalties, reducing the effective dimensionality of transport problems to obtain reasonable approximations of Wasserstein distances at lower computational costs. The annealing algorithm uses larger temperature parameters in early training stages to improve convergence speed, then gradually decreases them to obtain precise values.

To prevent different topic modeling results from being overly similar, a similarity loss is added to the topic composition matrix *t*:

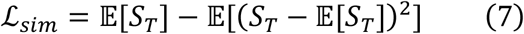

where 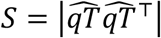 and 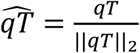 represents the cosine similarity between different topics. ℒ*_sim_* is designed as the mean minus variance of cosine values, encouraging uniform distribution of topic vectors. The final topic loss is a weighted sum of the prior gene program (PGP) optimal transport loss and similarity loss:

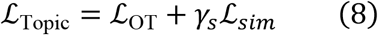

In the TopicVI model, distinctions are made among input PGPs to accommodate their biological meanings. Specifically, input PGPs are labeled as either cell type-related or background PGPs. For cell type-related PGPs, weighted transformations are applied in subsequent clustering loss calculations to emphasize the guidance of these clustering-related PGPs for cell representations (see Supplementary Text). Additionally, TopicVI can construct gene modules not guided by PGPs. In practice, we found that combining knowledge-guided module construction with partially knowledge-unguided module construction may improve overall model performance. Therefore, in subsequent experiments, we selected 25% of gene modules to be unguided by prior biological knowledge, with only knowledge-guided topics included in the optimal transport process.

### Deep Learning for Cell Clustering

The prediction of cell-cluster probabilities uses cell representations *z* output from the first module as input. This deep clustering model is constructed based on the DCE framework ^68^, with core features including: multiple differentiable clustering center parameters *μ*; adoption of differentiable K-means variants to optimize clustering center positions; and specific clustering loss functions. The default clustering process based on the Leiden algorithm ^69^ is performed to initialize parameters *μ*.

For any cell *i*, its probability of belonging to cell cluster *j* is calculated as follows:

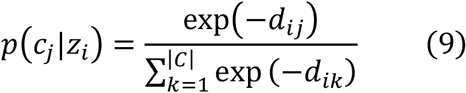

where *d*_*ij*_ = ||*z*_*i*_ − *μ*_*j*_||_2_ represents the Euclidean distance between sample embedding representation *z*_*i*_ and clustering center *μ*_*j*_. To adjust parameter *μ* during training, the clustering loss consists of three components: adaptive loss, deep clustering loss, and cluster centrality loss. The adaptive loss is defined as follows:

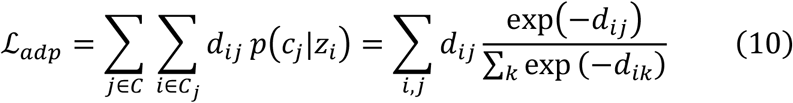

The main function of adaptive loss is to reasonably disperse samples and prevent cell cluster collapse.

The deep clustering loss is based on the DCE model ^68^, which adopts a self-training framework and defines the observed distribution *q* as:

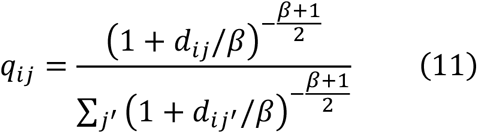

where *β* is a hyperparameter, set to 10 by default in this study. The target distribution *p* is defined as:

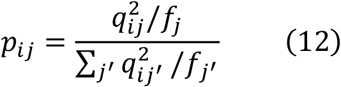

where *f*_*j*_ = ∑_*i*_ *q*_*ij*_ represents clustering frequency. The DCE loss is defined as the Kullback-Leibler (KL) divergence between them:

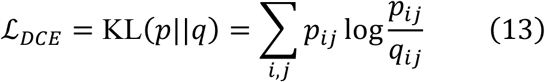

The cluster centrality loss function is defined similarly to the Davies-Bouldin index, used to evaluate intra-cluster distances and inter-cluster distances:

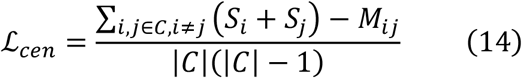

where 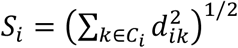 represents the sum of intra-cluster distances; *M* = ||*μ* − *μ*_*j*_||_2_ represents the distance between clustering centers; |*Q*| represents the number of cell clusters.

The total clustering loss is a weighted sum of the above three losses, with weights determined according to loss magnitudes:

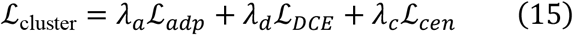

The complete loss function can be expressed as:

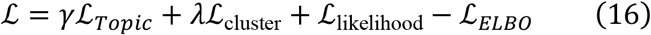

where the hyperparameters include **H** = {*γ, λ, γ*_*s*_, *λ*_*a*_, *λ*_*a*_, *λ*_*c*_}.

### Topic Analysis Based on TopicVI Model

To determine the genes in each topic, we scanned the matrix 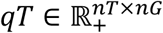 for non-zero entries, defining a topic as *t* = {*g*|*qT*_*t,n*_ ≠ 0}. To determine the related biological function of a topic, the Jaccard index is used to measure similarity between the prior topic and target topic. The Jaccard index is defined as: *J*(*t, t*_*P*_) = |*t* ∩ *t*_*P*_|/|*t* ∪ *t*_*P*_|. To identify the most relevant topics for each cluster, we use Student’s t-test to find differentially expressed topics across clusters based on topic loadings with Benjamini-Hochberg adjustment. This process is analogous to finding differentially expressed genes in scRNA-seq analysis and is implemented using “scanpy.tl.rank_genes_groups”.

## Data and Code Availability

The PBMC10k dataset was collected and processed using the scVI Python package ^24^ and can be accessed via the code `scvi.data.pbmc_dataset()`. The PBMC68k dataset was originally generated by Zheng et al. ^35^ and processed by Tamim et al. ^34^. The filtered and preprocessed h5ad file is available for download from Zenodo (https://zenodo.org/records/3357167). The Human Lung Cell Atlas (HLCA) ^26^ core data is publicly available through cellxgene (https://cellxgene.cziscience.com/collections/6f6d381a-7701-4781-935c-db10d30de293). The drug-perturbed GBM data (ZhaoSims2021) was originally generated by Zhao et al. ^41^ and subsequently processed by the scPerturb project ^70^, available at https://zenodo.org/record/7041849/files/ZhaoSims2021.h5ad. The spatial transcriptome dataset of human frontal cortex ^9^ is deposited in the NCBI Gene Expression Omnibus database under accession GSE233208. The preprocessed TCGA-GBM bulk RNA sequencing data and curated clinical metadata were obtained from the UCSC Xena browser.

The source code for TopicVI and the code for reproducing all analyses presented in this study is available on GitHub (https://github.com/zjupgx/topicvi and https://github.com/zjupgx/TopicVI-reproduce).

