## Supplementary Figure for "TopicVI: A Knowledge-guided deep interpretable model for resolving context-specific gene programs"

**A**

| Method | Topic Performance | | | | Bio conservation | | | | | Batch correction | | | | | Aggregate |
|---|---|---|---|---|---|---|---|---|---|---|---|---|---|---|---|
| | Module Coherence | Module Diversity | Gene-topic coherence | Topic Significance | Isolated labels | KMeans NMI | KMeans ARI | Silhouette label | cLISI | Silhouette batch | iLISI | KBET | Graph connectivity | PCR comparison | Total |
| **TopicVI** | 0.29 | 0.61 | 0.49 | 0.74 | 0.73 | 0.88 | 0.83 | 0.78 | 0.99 | 0.68 | 0.39 | 0.48 | 0.91 | 0.86 | 0.67 |
| **TopicVI (denovo)** | 0.24 | 0.97 | 0.46 | 0.80 | 0.64 | 0.71 | 0.57 | 0.82 | 0.97 | 0.48 | 0.16 | 0.26 | 0.86 | 0.94 | 0.63 |
| **scHPF** | 0.78 | 0.84 | 0.54 | 0.29 | 0.83 | 0.74 | 0.69 | 0.83 | 0.92 | 0.57 | 0.11 | 0.21 | 0.93 | 0.11 | 0.57 |
| **expiMap** | 0.48 | 0.05 | 0.00 | 0.45 | 0.52 | 0.80 | 0.73 | 0.53 | 0.67 | 1.00 | 0.68 | 0.66 | 0.49 | 0.95 | 0.49 |
| **LDVAE** | 0.34 | 0.51 | 0.19 | 0.00 | 0.77 | 0.83 | 0.82 | 0.79 | 0.98 | 0.67 | 0.10 | 0.19 | 0.95 | 0.29 | 0.46 |
| **Spike-slab LDA** | 0.02 | 0.33 | 0.82 | 0.29 | 0.66 | 0.45 | 0.44 | 0.54 | 0.74 | 0.39 | 0.36 | 0.24 | 0.78 | 0.05 | 0.42 |
| **Spectra** | 0.18 | 0.58 | 0.68 | 0.73 | 0.41 | 0.30 | 0.11 | 0.29 | 0.81 | 0.23 | 0.32 | 0.31 | 0.80 | 0.06 | 0.42 |
| **DcjComm** | 0.85 | 0.93 | 0.22 | 0.95 | 0.15 | 0.11 | 0.06 | 0.23 | 0.69 | 0.09 | 0.38 | 0.24 | 0.53 | 0.69 | 0.41 |
| **LIGER** | 0.98 | 0.82 | 0.21 | 0.40 | 0.39 | 0.03 | 0.04 | 0.24 | 0.10 | 0.47 | 0.98 | 0.80 | 0.23 | 1.00 | 0.41 |
| **Amortized LDA** | 0.05 | 0.04 | 0.91 | 0.08 | 0.46 | 0.32 | 0.26 | 0.48 | 0.75 | 0.46 | 0.26 | 0.25 | 0.80 | 0.16 | 0.36 |
| **Tree-spike-slab LDA** | 0.01 | 0.09 | 0.84 | 0.22 | 0.61 | 0.40 | 0.37 | 0.39 | 0.67 | 0.27 | 0.40 | 0.24 | 0.70 | 0.02 | 0.36 |
| **MuVI** | 0.36 | 0.03 | 0.11 | 0.00 | 0.23 | 0.29 | 0.10 | 0.29 | 0.88 | 0.54 | 0.09 | 0.15 | 0.17 | 0.56 | 0.24 |
| **CoGAPS** | 0.33 | 0.34 | 0.14 | 0.21 | 0.21 | 0.00 | 0.00 | 0.12 | 0.00 | 0.14 | 1.00 | 0.86 | 0.16 | 1.00 | 0.22 |

**Figure S1. Benchmarking methods across three primary evaluation domains.** Detailed sub-metric performance comparison across 14 evaluation criteria. The total score is calculated as the geometric mean of three domain-level scores: topic performance, biological conservation, and batch correction. Each domain score represents the mean of its corresponding sub-metrics.

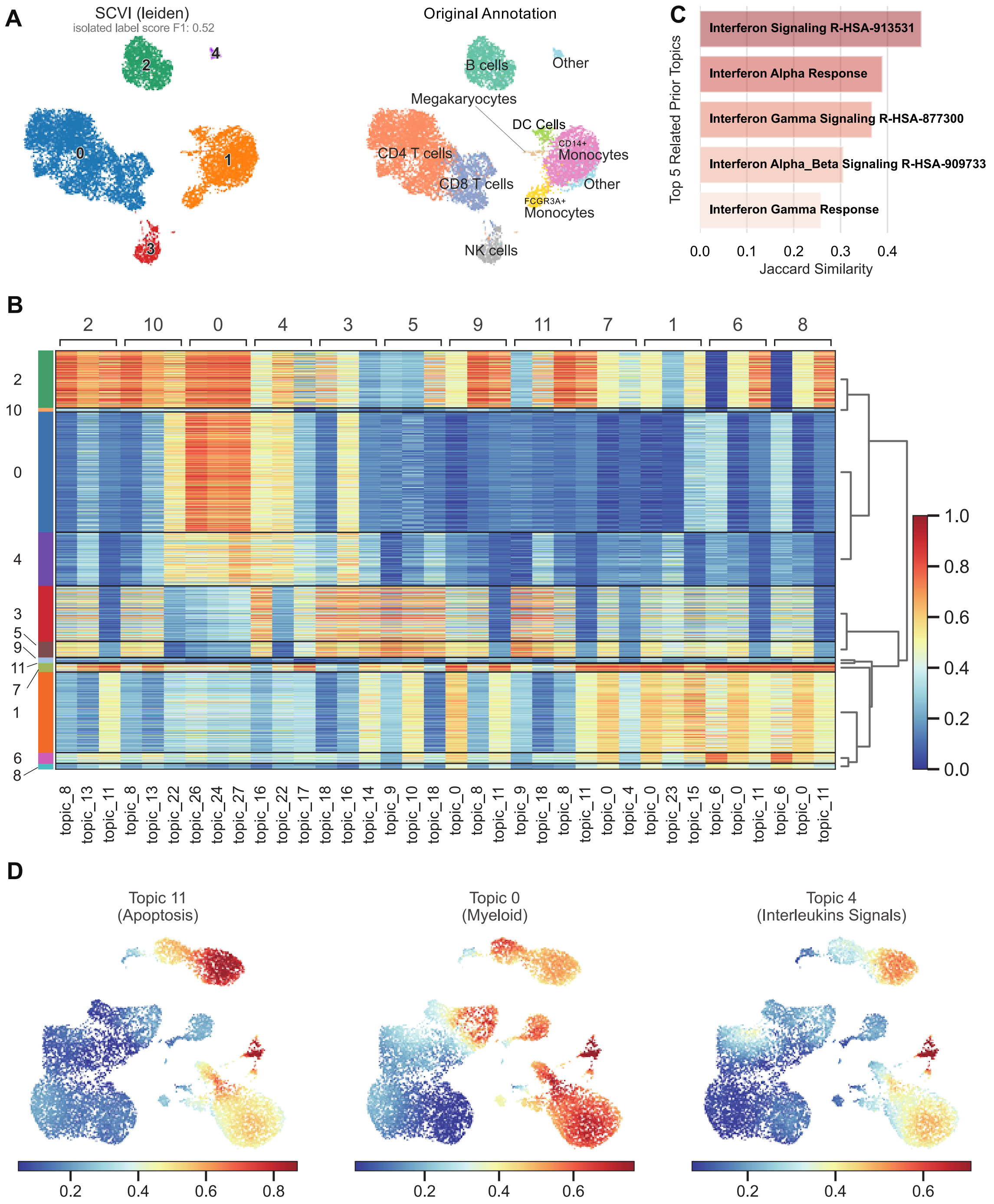


**Figure S2. Topic modeling enhances cell type annotation beyond standard SCVI modeling in PBMC10k dataset.** (A) SCVI model representation of the PBMC10k dataset. The left panel displays cell embeddings with Leiden clustering labels, while the right panel shows the original cell type annotations. SCVI fails to distinguish the subtypes in T cells and myeloid cells. (B) Student's t-test was used to identify cluster-specific differential loading topics, analogous to differential gene expression analysis. (C) Topic loadings and their corresponding annotations facilitate the distinction of cell clusters with different functional characteristics. Topic 11 (apoptosis-related), Topic 0 (myeloid cell markers), and Topic 4 (interleukin signaling) collectively enable the annotation of cluster 7 as neutrophils or macrophages.

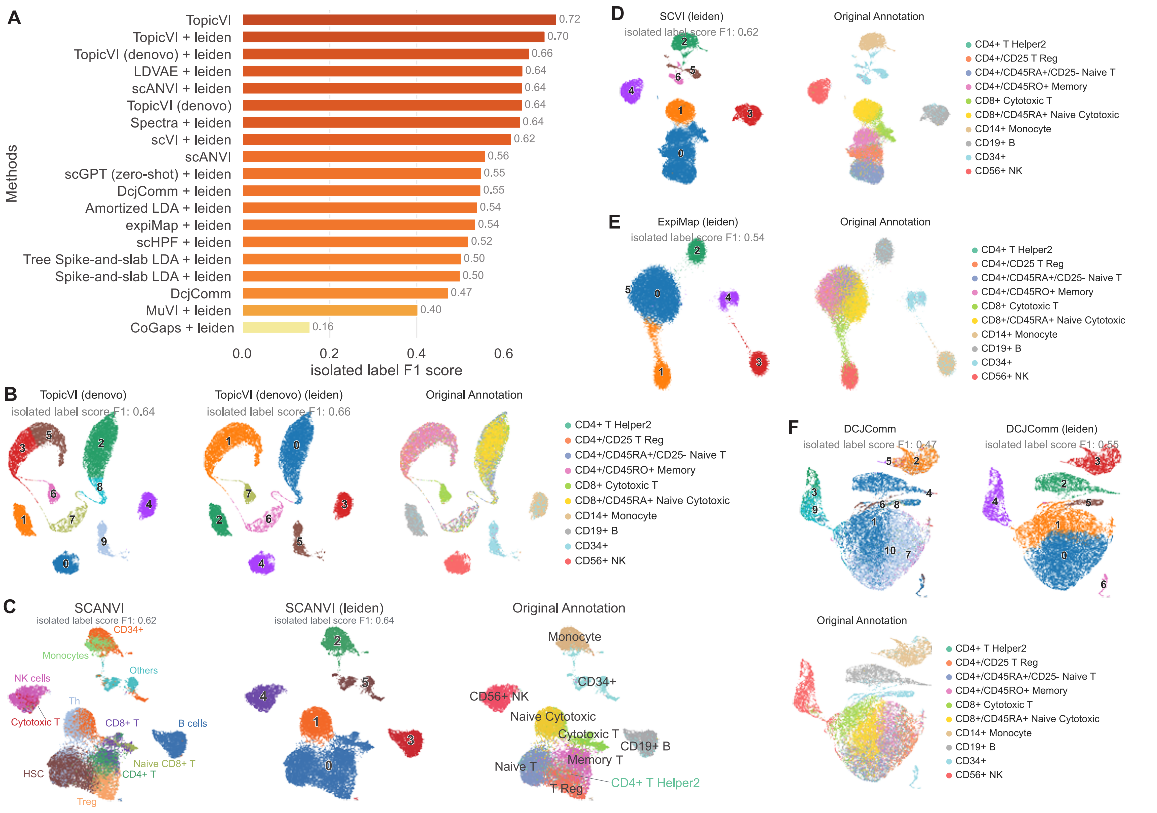


**Figure S3. Benchmark comparison reveals limitations of existing models in T cell subtype identification.** (A) Isolated label F1 scores for all evaluated methods. Leiden clustering was also applied to model-derived embeddings for comparison. (B-F) some of results of methods are displayed by UMAP. Representative UMAP visualizations from selected methods. TopicVI (denovo), scANVI, and DcjComm offer intrinsic cell type prediction and display predicted labels, Leiden clustering results, and original annotations. In contrast, ExpiMap and scVI lack built-in cell type prediction; their results include Leiden clustering outputs alongside manually curated reference annotations.

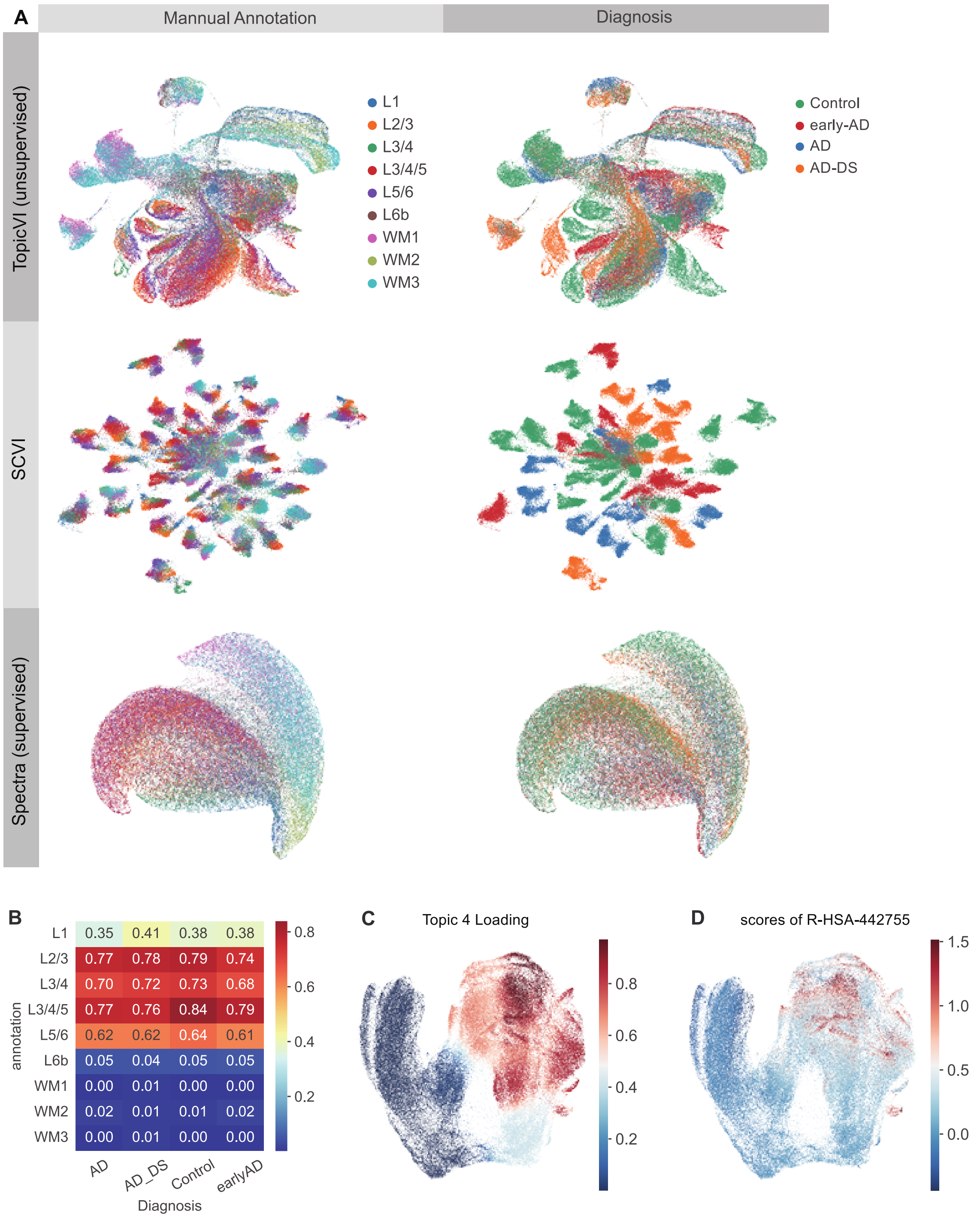
A
Mannual Annotation
Diagnosis
TopicVI (unsupervised)
SCVI
Spectra (supervised)
L1
L2/3
L3/4
L3/4/5
L5/6
L6b
WM1
WM2
WM3
Control
early-AD
AD
AD-DS
B
L1 0.35 0.41 0.38 0.38
L2/3 0.77 0.78 0.79 0.74
L3/4 0.70 0.72 0.73 0.68
L3/4/5 0.77 0.76 0.84 0.79
L5/6 0.62 0.62 0.64 0.61
L6b 0.05 0.04 0.05 0.05
WM1 0.00 0.01 0.00 0.00
WM2 0.02 0.01 0.01 0.02
WM3 0.00 0.01 0.00 0.00
annotation
AD
AD_DS
Control
earlyAD
Diagnosis
0.8
0.6
0.4
0.2
C
Topic 4 Loading
0.8
0.6
0.4
0.2
D
scores of R-HSA-442755
1.5
1.0
0.5
0.0

**Figure S4. Topic-based approach shows superior discrimination of cell clusters compared to pathway analysis.** (A)**(A)** Under unsupervised learning, cell clustering and representation are primarily driven by diagnostic status. Spectra (supervised) also struggles to distinguish anatomical annotations. The two UMAP panels show manual cell type annotations and disease diagnosis, respectively. (B) Annotation labels used in supervised learning guide Topic 4 loading patterns, which align closely with the chosen annotation type. (C-D) Comparison of Topic 4 loading (C) and gene set scores for Reactome pathway R-HSA-442755 (D). Topic 4 loading exhibits significantly greater discriminative power across diagnostic groups than the corresponding pathway score.

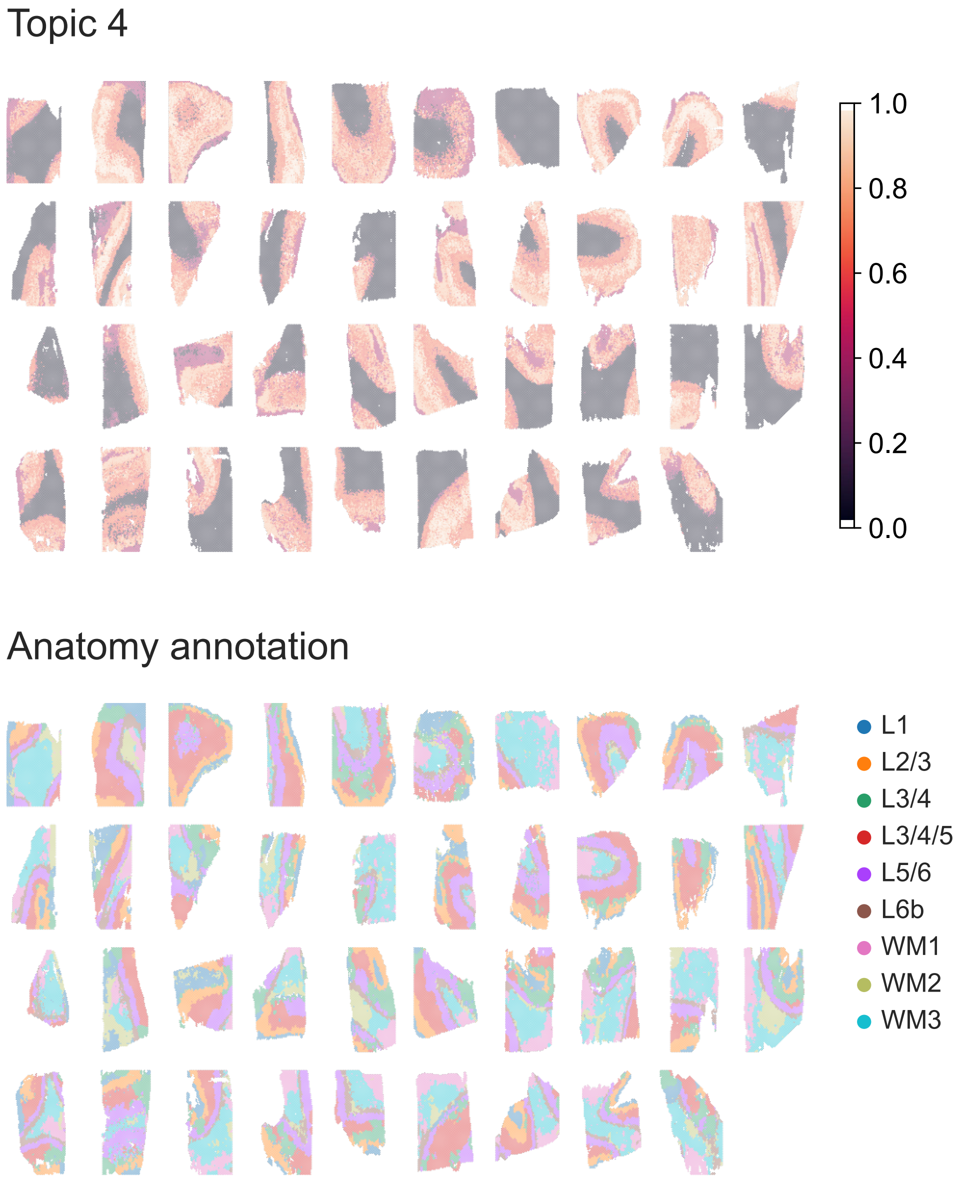


**Figure S5. Spatial validation of Topic 4 loading reveals cortical layer organization in Visium data.** Spatial distribution of Topic 4 loading (top) compared to manual anatomical annotations (bottom) across 39 brain tissue slices using 10x Visium spatial transcriptomics. Topic 4 loading effectively distinguishes cortical layers L1-L6 and captures spatial expression patterns that correspond to cortical laminar architecture, validating the biological relevance of the NMDA receptor-related topic modeling approach in spatial context.

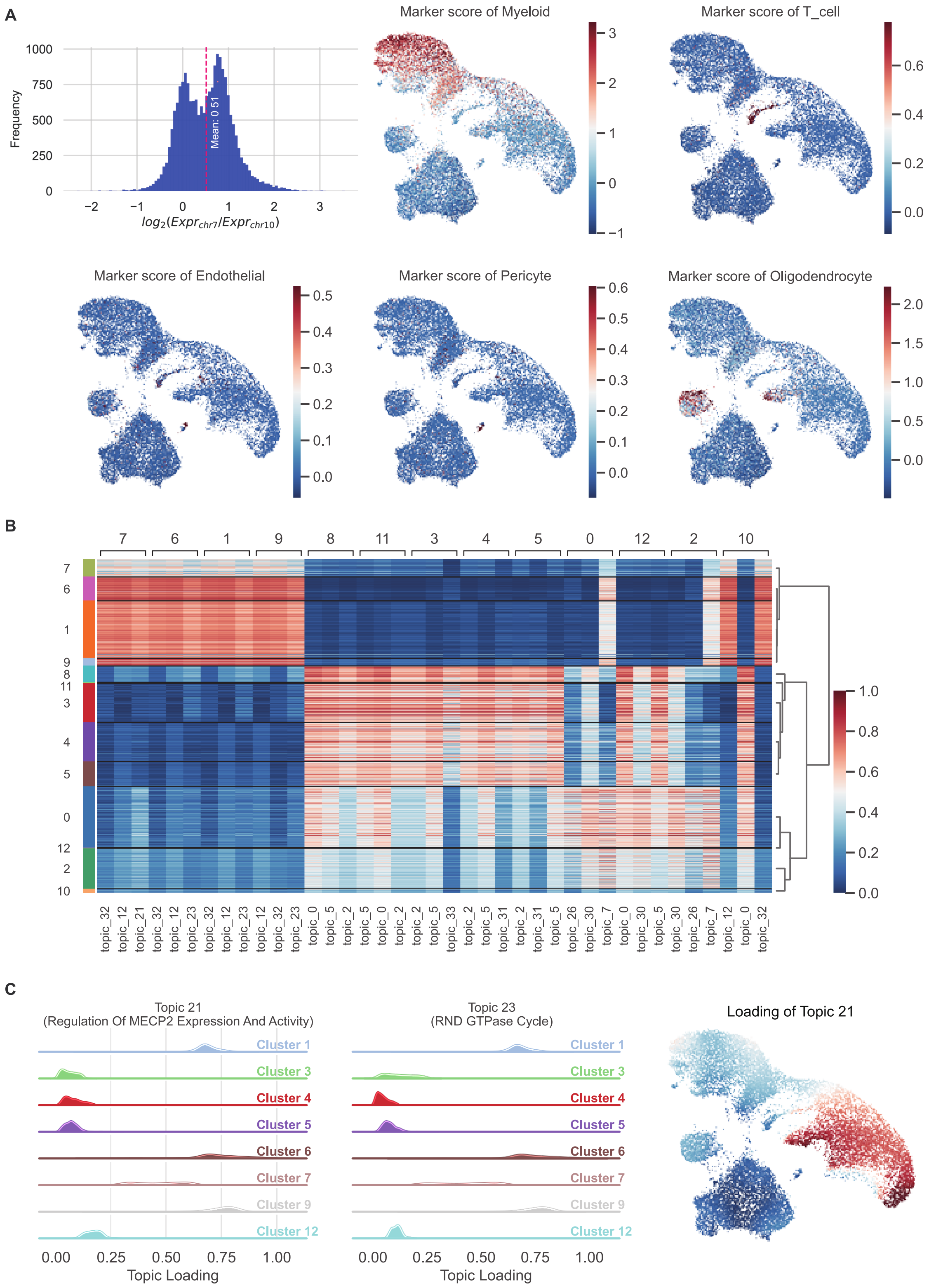

A
Frequency
Mean: 0.51
$log_2(Expr_{chr7}/Expr_{chr10})$
Marker score of Myeloid
Marker score of T_cell
Marker score of Endothelial
Marker score of Pericyte
Marker score of Oligodendrocyte
B
C
Topic 21
(Regulation Of MECP2 Expression And Activity)
Topic 23
(RND GTPase Cycle)
Cluster 1
Cluster 3
Cluster 4
Cluster 5
Cluster 6
Cluster 7
Cluster 9
Cluster 12
Topic Loading
Loading of Topic 21

**Figure S6. TopicVI identifies drug response signatures and malignancy patterns in GBM cells.** (A) Transfer of cell type marker scores and malignancy signatures. The malignancy score is defined as the log-transformed ratio of average expression between chromosome 7 and chromosome 10, reflecting Chr. 7 amplification and Chr. 10 deletion characteristic of GBM malignancy. Cell type markers were derived from the original research publication. (B) Student's *t*-test analysis identifying cluster-specific topics from 40 total topics. (C) Topic distribution across cell clusters. Topic 21 corresponds to regulation of MECP2 expression and activation pathways, showing elevated expression in mesenchymal glioblastoma (GBM) subtypes. Topic 23 is associated with RND GTPase cycle processes, where RND1 functions as a novel p53 controller and positive regulator of p53 signaling, promoting ferroptosis in GBM.

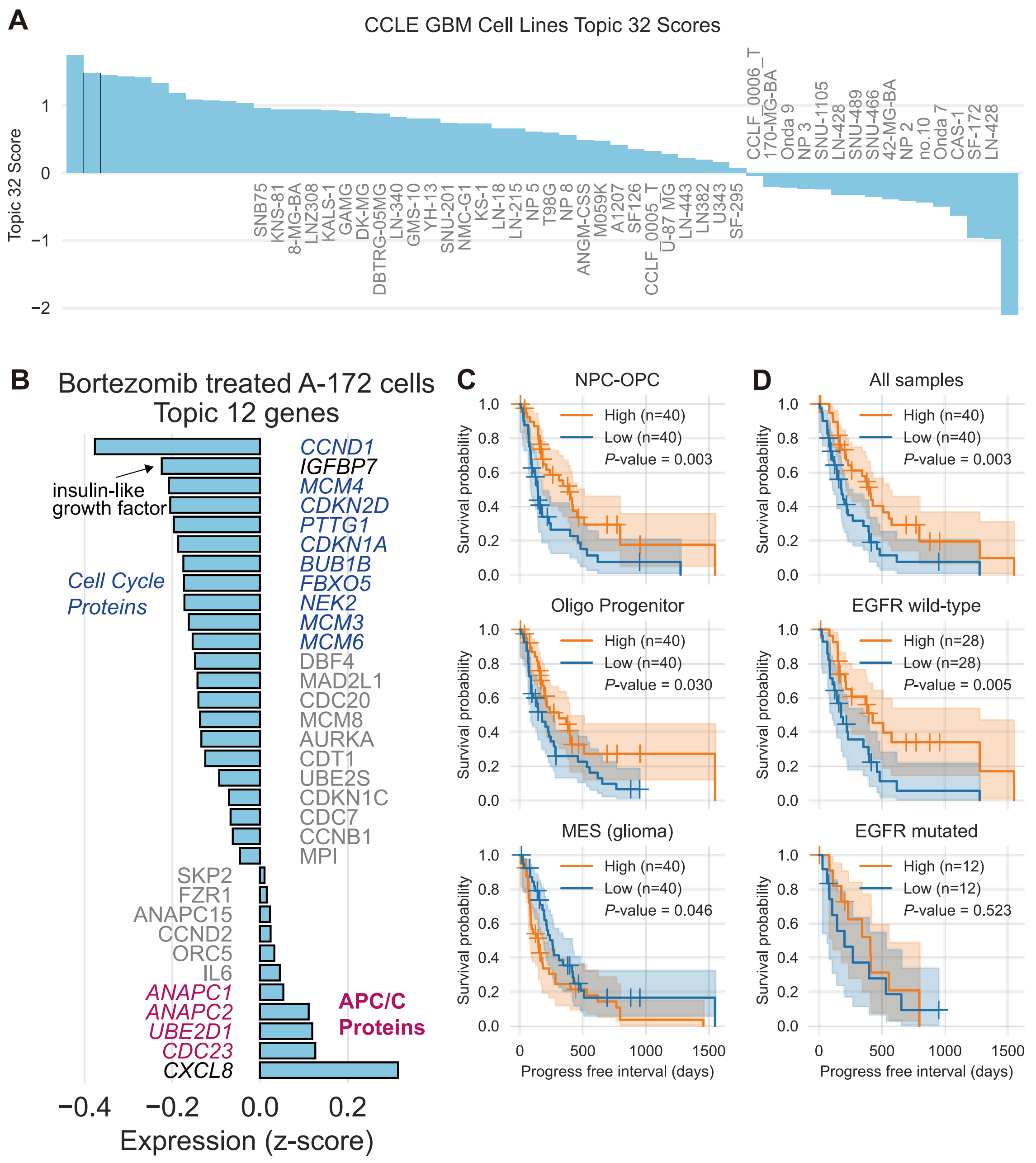


**Figure S7. Survival Associations and Drug Sensitivity Patterns of Topics in GBM Cell Lines.** (A) Topic 32 scores across CCLE GBM cell lines. The A-172 cell line ranks among the top two in Topic 32 score. (B) Expression of Topic 12 genes (z-transformed) in Bortezomib-treated A-172 cells. Cell cycle-related proteins are labeled in blue, APC/C complex proteins in red, and genes with minimal expression change (near zero) in grey. (C) Three glioblastoma (GBM)-related metaprograms show significant associations with patient survival (log-rank test). (D) Kaplan–Meier analysis of relationship between progress free interval and Topic 32 score in three cohorts: all samples in TCGA-GBM, *EGFR* mutated individuals and *EGFR*-wild type individuals (log-rank test).
