## Supplementary Text for "TopicVI: A Knowledge-guided deep interpretable model for resolving context-specific gene programs"

### Details of TopicVI model

#### Evidence Lower Bound

TopicVI employs a Bayesian modeling framework to model gene expression data $X\in\mathbb{R}^{nc\times ng}$, where $ng$ represents the number of genes in gene set $\mathbb{G}$ and $nc$ represents the number of cells in cell set $\mathbb{C}$. The model assumes that cells can be partitioned into several cell clusters $C_{i}\mathbb{\subset C}$, while gene sets may belong to multiple prior feature modules or topics $G_{i}\mathbb{\subset G}$. It is noteworthy that gene topics may not contain all gene sets, but cell types must cover all cells, satisfying $\bigcup_{i}C_{i}\mathbb{=C}$.

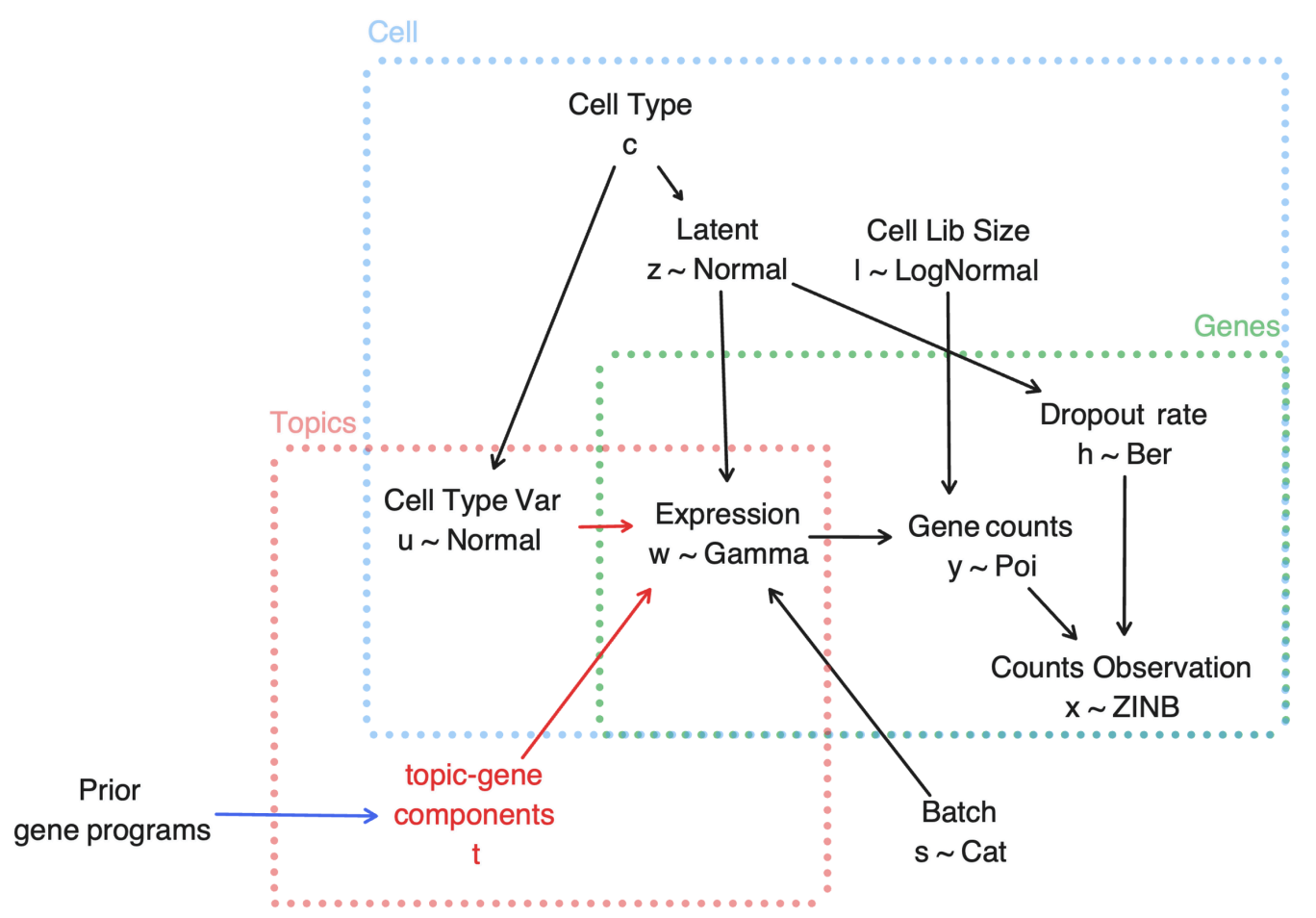

The generation process of gene expression is illustrated in the accompanying figure. Specifically, this study assumes that the variational posterior distribution for gene expression $x=X_{i}$ of a single cell can be factorized as:

$$q(c,z,u,l|x,t_{P})=q(z|x)q(c|z)q(u|z,c)q(l|x) (1)$$

where the variables are defined as follows: $x$ is the cell expression vector; $c$ is the cell type label; $z$ is the cell embedding representation; $u$ is the feature representation specific to cell types; $l$ is the cell sequencing depth (library size); and $t_{P}$ represents prior information.

For sequencing depth, the library size factor is calculated based on the following formula:

$$l=\frac{\text{cell\_total}}{\exp\left( \frac{1}{n}\sum_{i=1}^{n} \log(\text{cell\_total}_{i}) \right)} (2)$$

where $\text{cell\_total}_{i}=\sum_{j} X_{ij}$ represents the total expression of cell $i$. Thus, the library size factor is the total cell set expression divided by the geometric mean across all cell sets.

According to eq. 1, the loss function for maximizing the Evidence Lower Bound (ELBO) can be derived:

$$\log p(x,t_{P})\geq\mathbb{E}_{q(c,z,u,l|x,t_{P})}[logp(x,z,u,c,l)-logq(c,z,u,l|x,t_{P})] (3)$$

Expanding eq. 3, the ELBO loss function consists of the following components:

$$\begin{aligned} \mathcal{L}_{ELBO}&=\underset{\text{reconstruction loss}}{\underbrace{\mathbb{E}_{q(z,u,c,l|x,t_{P})}[logp(x|z,l)]}} \\ &+\underset{\text{conditional generation loss}}{\underbrace{\mathbb{E}_{q(z,u,c,l|x,t_{P})}[logp(z|u,c)-logq(z|x)]}} \\ &+\underset{\text{clustering prior loss}}{\underbrace{\mathbb{E}_{q(z,u,c,l|x,t_{P})}[logp(c)-logq(c|z)]}} \\ &+\underset{\text{u prior loss}}{\underbrace{\mathbb{E}_{q(z,u,c,l|x,t_{P})}[logp(u)-logq(u|z,c)]}} \\ &+\underset{\text{l prior loss}}{\underbrace{\mathbb{E}_{q(z,u,c,l|x,t_{P})}[logp(l)-logq(l|x)]}} \end{aligned} (4)$$

**1) Reconstruction Loss**

$$\mathbb{E}_{q(z,u,c,l|x,t_{P})}[logp(x|z,l)]=\mathbb{E}_{q(z|x)q(l|x)}[logp(x|z,l)] (5)$$

**2) Conditional Generation Loss**

$$\begin{aligned} \mathbb{E}_{q(z,u,c,l|x,t_{P})}&[logp(z|u,c)-logq(z|x)]= \\ &\mathbb{E}_{q(z|x,t_{P})}[\sum_{c=1}^{K} q(c|z)\mathbb{E}_{q(u|z,c)}[logp(z|u,c)-logq(z|x)]] \\ +&\mathbb{E}_{q(z|x,t_{P})q(u|z,c)}[logp(z|u)] \end{aligned} (6)$$

**3) Prior Loss** The prior loss uses KL divergence to measure the difference between prior and posterior distributions, including: clustering $c$ (with uniform prior distribution); sequencing depth $l$ (with normal prior distribution calculated from expression levels); and cell type specificity $u$ (with standard normal prior distribution):

$$\left\{ \begin{matrix} \mathbb{E}_{q(z,u,c,l\mid x,P)}[logp(l)-logq(l\mid x)]=-KL(q(l\mid x)\parallel p(l)) \\ \mathbb{E}_{q\left( z,u,c,l\mid x,t_{P} \right)}[logp(c)-logq(c\mid z)]=\mathbb{E}_{q(z\mid x)}[-KL(q(c\mid z)\parallel p(c))] \\ \mathbb{E}_{q\left( z,u,c,l\mid x,t_{P} \right)}[logp(u)-q(u\mid z,c)]=\mathbb{E}_{q\left( z\mid x,t_{P} \right)}\left[ \sum_{c=1}^{K} -q(c\mid z)KL(q(u\mid z,c)\parallel p(u)) \right] \end{matrix} \right. (7)$$

#### Hyperparameters

For the weights of different terms in the loss function, this research fixes the following hyperparameters: $\gamma_{s}=1000$, $\lambda_{a}=200$, and $\lambda_{d}=50$. The parameter $\gamma$ is determined based on the number of topics and priors according to the formula $\gamma=100\log_{10}(n_{T}\times n_{TP}+1)$, where $n_{T}$ is the number of topics (default value of 32), and $n_{TP}$ is the number of prior topics. The clustering weights $\lambda_{c}$ and $\lambda$ are set empirically. For specific parameter settings, please refer to the source code available on GitHub.

In our experience, most hyperparameters do not significantly influence TopicVI results, with the main consideration being the ‘clustering strength’, which depends on data diversity and the degree of cluster separability. Another critical hyperparameter that must be determined is the number of clusters. For most conditions, if the exact number of clusters is unknown, we recommend using a relatively large number, as clusters may collapse by pushing some cluster centers away from most cells during optimization.

#### Alternative Model Choices

##### Training Mode

The default training mode of TopicVI is unsupervised. For conditions where clustering is difficult or cell type labels are known, especially when there are many confounding sources or clustering based on cell transcriptomes is insufficient, we have also implemented a supervised training mode for TopicVI. Similar to scANVI’s semi-supervised training ^1^, a classification loss is included in the supervised learning process. We use confidence-modulated cross-entropy loss to guide the classification task:

$$\mathcal{L}_{\text{classification}}=\frac{1}{N}\sum_{i=1}^{N} \left( 1-p(c_{y_{i}}\mid z_{i}) \right)^{2}\cdot\left( -logp(c_{y_{i}}\mid z_{i}) \right) (8)$$

where $c_{y_{i}}$ is the provided label of sample $i$. This focal loss ^2^ has the advantage of: (1) penalizing samples for which the model assigns low probability to the correct class, (2) suppressing easy examples by reducing the loss for high-confidence predictions, and (3) encouraging the model to focus on harder examples, which is particularly useful for addressing class imbalance or noisy labels.

##### Prior Selection

As mentioned in the Methods section of the main manuscript, there are two types of prior gene programs (PGP) considered in TopicVI: cell type markers and background priors. The key difference lies in clustering, where a weighted embedding distance is calculated for the cluster loss. For cell embedding $z_{i}\in\mathbb{R}^{1\times k}$ of cell set $i$ and cluster center $\mu_{j}$, the weighted distance $\hat{d}_{ij}$ is defined as:

$$\hat{d}_{ij}=\sqrt{\sum_{m=1}^{k} w_{m}\cdot(z_{im}-\mu_{jm})^{2}} (9)$$

where $w$ is the weight of latent dimensions, calculated by the tensor product of model parameters:

$$w_{m}=\frac{1}{nG}\sum_{i=1}^{nG} \text{softmax}\left( \sum_{j=1}^{nT_{Pc}} \mu_{jm}\cdot\mathcal{T}_{ij} \right) (10)$$

where $\mathcal{T=}\text{softmax}(qT_{[T_{Pc},:]})$, and $T_{Pc}$ represents the prior gene sets that are assigned as cell type markers or cluster-aware gene sets.

### Details of Benchmarking

#### Building Standard Datasets

Based on previous research ^3^, this study conducted stratified sampling from the core data of the Human Lung Cell Atlas (HLCA) ^4^ according to annotation hierarchies, constructing a benchmark dataset comprising 8 subsets. It is particularly noteworthy that the original HLCA manual annotation system contains five hierarchical levels of labels, with annotation depth correlating with the degree of cell type subdivision.

Among the 8 constructed data subsets, 4 employed balanced sampling strategies (with equal numbers of cell sets across subtypes), while 4 adopted unbalanced sampling strategies (maintaining original cell type distribution ratios). The stratified sampling was performed based on the hierarchical cell type labels, ensuring representative coverage across different annotation levels while preserving the biological diversity within each cell type category. Through this design, the dataset effectively simulates single-cell data characteristics of varying complexity levels encountered in real-world scenarios. Detailed characteristics of each dataset subset are provided in Supplementary Table 2.

For each dataset’s characteristics, this study selected appropriate cell type marker genes from the aforementioned pre-collected prior gene programs (PGP) based on the original cell labels, searching for cell markers in CellMarkerDB V2 ^5^and scType ^6^. All datasets’ background gene programs integrate both MSigDB Hallmark ^7^ and Reactome pathways ^8^.

#### Methods in Benchmarking

The methods evaluated in this study are primarily categorized into three types: VAE-based models, NMF-based methods, and pre-trained large models.

**VAE-based models:** SCVI ^9^ serves as the first framework to apply VAE for single-cell transcriptomic batch correction and integration, establishing the foundation for subsequent research. SCANVI ^1^ builds upon SCVI by introducing cell type label inputs and employing semi-supervised learning strategies to enhance predictive capabilities for new data. Its implementation follows the seed labeling strategy recommended in the official documentation, which first constructs high-confidence labels based on overall expression levels, then proceeds with semi-supervised learning. ExpiMap ^10^ modifies SCVI’s multilayer perceptron by designing gene program selectors to optimize gene program construction. The LDVAE model ^11^ replaces neural networks with linear functions to achieve precise gene weighting, enabling each embedding dimension to characterize specific gene programs. Amortized LDA ^12^ integrates topic modeling into the VAE framework, utilizing cell set embeddings to infer gene topic composition. Tree-spike-slab LDA ^13^ and Spike-slab LDA ^14^ introduce spike-slab prior assumptions based on LDA, positing that gene weights within gene programs exhibit sparse distribution characteristics, with Tree-spike-slab additionally assuming hierarchical tree-like relationships among topics.

**NMF-based methods:** NMF-based methods construct models through decomposition of gene expression, with decomposition results containing cell set loadings on gene programs and weight distributions of genes within gene programs. ScHPF ^15^ constructs a hierarchical Poisson decomposition framework to accommodate single-cell data characteristics. MuVI approaches the problem from the gene perspective, integrating prior gene sets ^16^ into the decomposition process, while DcJcomm ^17^ incorporates cell set clustering information into the decomposition process from the cellular perspective. Spectra ^18^ distinguishes between cell type-specific and non-specific gene programs during NMF decomposition. CoGAPS ^19^ and LIGER ^20^ focus on NMF decomposition in multi-dataset integration scenarios.

**Pre-trained large models and other methods:** Additionally, Harmony ^21^ achieves batch integration through projection in cell set embedding space and represents one of the most widely applied methods currently. Recently emerging single-cell transcriptomic pre-trained large models, such as scGPT ^22^, demonstrate powerful downstream task processing capabilities, including cell set representation construction and gene weight interpretation analysis.

The TopicVI method proposed in this study innovatively combines the advantages of VAE and NMF for topic modeling, comprising two core modules: cell embedding construction and a deep biclustering module. To investigate the impact of prior gene programs (PGP) on cell embedding representation and gene program construction, this study designed a control model “TopicVI (denovo)”, which excludes PGP input.

#### Evaluation Metrics

This study established a multidimensional evaluation system based on the scIB ^21^ assessment framework.

**Batch effect removal assessment:** The following metrics were employed: k-nearest-neighbor batch effect test (kBET ^23^) to quantify the degree of batch mixing; cross-batch k-nearest-neighbor graph connectivity to reflect data integration effectiveness; Average Silhouette Width (ASW ^23^) to evaluate batch separation degree; graph integration Local Inverse Simpson’s Index (graph iLISI ^21^) to measure local diversity; and PCA regression ^23^ to analyze residual batch effects in principal components.

**Bio-conservation evaluation:** The evaluation of bio-conservation was conducted from two dimensions: cell type label conservation and label-free conservation. Specifically, cell-type Local Inverse Simpson’s Index (cLISI ^21^) was employed to assess local neighborhood homogeneity; clustering label accuracy was quantified through Adjusted Rand Index (ARI) and Normalized Mutual Information (NMI); cell-type ASW evaluated inter-class separation degree; and isolated label scores ^21^ were used to evaluate the identification effectiveness of rare cell types.

**Topic assessment:** Topic assessment was based on the OCTIS ^24^ framework, encompassing four core metrics: (1) Module Coherence evaluates semantic consistency within topics by calculating mutual information between gene expression distributions and module distributions; (2) Module Diversity quantifies the degree of difference between topics using rank-biased overlap computation methods; (3) Gene Topic Coherence assesses the correlation between modules and gene expression using cosine similarity; and (4) Topic Significance evaluates the degree of difference between topic distributions and random distributions by calculating the average KL divergence between topic-gene distributions and three reference distributions (uniform distribution, null distribution, and background distribution) ^25,26^. During the evaluation process, this study selected genes with scores greater than 0 and ranked in the top 50 from topic-gene distributions for analysis.
